# Big Data in Myoelectric Control: Large Multi-User Models Enable Robust Zero-Shot EMG-based Discrete Gesture Recognition

**DOI:** 10.1101/2024.07.11.603119

**Authors:** Ethan Eddy, Evan Campbell, Scott Bateman, Erik Scheme

## Abstract

Myoelectric control, the use of electromyogram (EMG) signals generated during muscle contractions to control a system or device, is a promising modality for enabling always-available control of emerging ubiquitous computing applications. However, its widespread use has historically been limited by the need for user-specific machine learning models because of behavioural and physiological differences between users. Leveraging the publicly available 612-user EMG-EPN612 dataset, this work dispels this notion, showing that true zero-shot cross-user myoelectric control is achievable without user-specific training. By taking a discrete approach to classification (i.e., recognizing the entire dynamic gesture as a single event), a classification accuracy of 93.0% for six gestures was achieved on a set of 306 unseen users (who provided no training data), showing that big data approaches (compared to most EMG studies, which typically employ only 10-20 users) can enable robust cross-user myoelectric control. By organizing the results into a series of mini-studies, this work provides an in-depth analysis of discrete cross-user models to answer unknown questions and uncover new research directions. In particular, this work explores the number of participants required to build cross-user models, the impact of transfer learning for fine-tuning these models, and the effects of under-represented end-user demographics in the training data, among other issues. Additionally, in order to further evaluate the performance of the created cross-user models, a completely new data set was created (using the same recording device) that includes known covariate factors such as cross-day use and limb-position variability. The results show that the large data models can effectively generalize to new datasets and mitigate the impact of common confounding factors that have historically limited the adoption of EMG-based inputs.

## 1 INTRODUCTION

Modern-day human-computer interaction (HCI) is rapidly evolving. Devices have become smaller, processing power continues to increase, and technology is more interconnected than ever before, expanding the need for and possibility of interactions beyond traditional tethered desktop environments. This shifting paradigm towards ubiquitous computing (Friedewald and Raabe, 2011; Weiser, 1999), where the goal is to “[enhance] computer use by making many computers available throughout the physical environment, while making them effectively invisible to the user” (Weiser, 1993). First envisioned in the late 20th century, this concept is quickly becoming a reality with the continued emergence of hardware devices like heads-up mixed reality displays (Speicher et al., 2019), wearable technologies (Dunne et al., 2014), and through advancements in artificial intelligence (Li et al., 2020).

The increased prevalence of ubiquitous computing has created a need for new interaction techniques that are reliable, convenient, and always available. While many inputs exist and have been adopted for such use cases, their widespread and consistent use is currently limited by the challenges of daily life. For example, computer vision does not work well in the dark or when an individual’s hands are occluded from the camera (e.g., behind their back or in their pocket) (Oudah et al., 2020). Alternatively, speech-based inputs are conspicuous, lack social acceptability, and can be hampered by environmental noise (Li et al., 2014; Pandey et al., 2021). So, it follows that there is a need for an input modality that can work under these frequently experienced, everyday, real-world conditions. Myoelectric control, the control of a system using the electrical signals generated when muscles are contracted, has been identified as one promising solution but has yet to achieve the reliability and robustness required to be sufficiently usable and widely adopted Eddy et al. (2023c).

The first myoelectric control systems can be traced back to the 1940s when a robotic limb that could be controlled through muscle contractions was developed (Englehart and Hudgins, 2003). Enabled by leveraging the surface electromyogram (EMG) signal—a direct representation of the motor unit activations corresponding to the level of muscle activation at a given recording site (Farina et al., 2004)—the muscle signals created by the human body became an enabling input with various potential applications. Researchers quickly realized that these signals, when recorded from the residual limb of an amputee, could be leveraged to control powered prostheses. A key advantage of this approach was that it offered a control strategy that worked in most real-world scenarios. Although promising, early myoelectric control systems were not particularly robust or intuitive and were far from capable of replicating the dexterity of a functional limb.

In the following decades, myoelectric control slowly evolved, with its primary use case continuing to be the control of powered prostheses (Huang et al., 2024; Chan et al., 2000; Farina et al., 2014). A notable advancement in this history came in 2003 through the introduction of continuous pattern recognition-based control (Englehart and Hudgins, 2003). Instead of being limited to a single degree of freedom (Scheme and Englehart, 2011; Hargrove et al., 2017), pattern-recognition systems leverage multiple electrodes distributed across several muscle sites to enable the mapping of a wider set of gestures and device commands. With this approach, gesture predictions are generated from *short windows of data at fixed increments* (on the order of milliseconds) to continuously make micro-adjustments to the device’s position. This control strategy has now been made commercially available (Infinite, 2024; COAPT, 2024; Ottobock, 2024), marking a significant step forward in the field of prosthetics. Since then, myoelectric prosthesis control has continued to evolve and mature, such as through the simultaneous control of degrees of freedom (Hahne et al., 2014; Smith et al., 2014) and through augmentations of the human body, including targeted muscle reinnervation and osseointegration (Farina et al., 2023).

The history of success of myoelectric control in prosthetics ultimately led to uptake within the HCI community, where researchers began to explore the potential of EMG for enabling input for other (non-prosthetic) applications in the 2000s (Saponas et al., 2009, 2008, 2010). In particular, as opposed to the continuous control schemes used for prosthesis control, some HCI practitioners began to explore the recognition of gesture sequences (e.g., finger taps or snaps) that could be mapped to discrete inputs, as these aligned better with common inputs to a computer system like button clicks Saponas et al. (2009). Instead of continuously making predictions on static contractions, these so-called *discrete myoelectric control systems* make a single prediction at the end of a dynamic gesture (e.g., a swipe of the wrist), meaning it is a many-to-one mapping between windows of EMG data and predictions from the classifier (Eddy et al., 2023c; Labs et al., 2024).

Regardless of the control strategy, the general exploration of myoelectric control in HCI was further facilitated in 2014 through the introduction of the *Myo Armband* by Thalmic Labs, a commercially available and inexpensive surface-EMG wearable device (Rawat et al., 2016; Benalcázar et al., 2017). Due to the accessibility that this device offered, including built-in gesture recognition, there was a period of rapid exploration and excitement (Haque et al., 2015; Zadeh et al., 2018; Koskimäki et al., 2017; Mulling and Sathiyanarayanan, 2015; Dai et al., 2021). However, this initial interest eventually subsided, as the technology, at the time, lacked the precision and robustness required for real-world consumer use (Karolus et al., 2022; Eddy et al., 2023c). Nevertheless, while advancements in the space slowed down, commercial products continued to emerge (e.g., the Mudra (Mudra, 2024) and Pison (Pison, 2024) devices), albeit none of which have yet become widely adopted.

A common factor shared between the advancement of myoelectric control in both communities (HCI and prosthetics) is that it has primarily occurred through publicly funded research in academic settings. Control schemes have been historically tested on a limited number of participants (e.g., *N <* 20*users*), and datasets are frequently recorded with unique hardware and seldom publicly released (Eddy et al., 2023c), slowing the progress of the field. Additionally, most EMG research in both communities has historically employed user-dependent models, wherein the model that deciphers user intent is trained and tested based on a single user’s data. While this may make sense for prosthesis control, where every amputee’s residual musculature differs, this is a fundamental limitation for general-purpose HCI applications, where consumer devices should be as close to “plug-and-play” as possible. However, plug-and-play convenience has yet to be fully realized, and users still largely have to undergo arduous and tedious data collection every time they put on an EMG device. Additionally, if not intentionally included in the training procedure, the resulting models lack robustness to confounding factors such as cross-day use, limb-position variation, and electrode shift, thus further exacerbating online usability issues (Campbell et al., 2020). While researchers have been able to alleviate some of these factors through transfer learning (Campbell et al., 2021; Jiang et al., 2024; Xu et al., 2024; Côté-Allard et al., 2019) and domain adaptation (Zhang et al., 2022; Eddy et al., 2023a; Campbell et al., 2024), these strategies fall short from the ideal zero-shot case whereby no training data from the end user is required, such as is the case for current computer-vision based gesture recognition systems (Oudah et al., 2020).

In 2019, the largest investment in the history of myoelectric control was made when Meta (i.e., Facebook) acquired Ctrl Labs (the company then owning the intellectual property of the Myo Armband) for reports of between $500 million to $1 billion (Rodriguez, 2019; Statt, 2019). This came around the same time that Ctrl Labs first revealed their technology to the research community, enabling *neuromuscular control* (i.e., myoelectric control) (Melcer et al., 2018). Recently, after nearly five years of closed-door development, Ctrl Labs (now Meta) has claimed that zero-shot cross-user myoelectric control is possible for a variety of applications, including (1) *1D cursor control*, (2) *discrete control through thumb swipes* and (3) *continuous handwriting* (Labs et al., 2024). Unlike the majority of EMG research, this work was conducted by a mega-cap multi-national company, a team of *∼*200 individual contributors, a closed-source 16-channel EMG armband worn on the wrist, and data collections consisting of thousands of participants. To the best of our knowledge, this work represents a massive step forward in the history of myoelectric control, leaving the research community in an exciting position.

### 1.1 Scope and Contribution

Despite recent demonstrations suggesting that cross-user models perform reasonably well with large amounts of data (Labs et al., 2024; Vásconez et al., 2023; Valdivieso Caraguay et al., 2023; Barona López et al., 2024), many questions remain unanswered. This work thus explores cross-user myoelectric control practices for discrete event-driven systems to uncover and guide future research directions and highlight best practices moving forward. Leveraging the publicly available EMG-EPN612 dataset (Benalcazar et al., 2021), the results of this work corroborate and expand on recent reports (Labs et al., 2024), showing that robust zero-shot discrete cross-user myoelectric control models that require no training from the end user are, in fact, possible. While this has been pursued over recent years (Kim et al., 2019; Lin et al., 2023), these works have been largely conducted on small datasets (*<*50 participants) compared to the thousands collected in (Labs et al., 2024) and the 612 participants (here called subjects) in (Benalcazar et al., 2021). Correspondingly, this work provides a commentary and analysis of the impact and role of “big data” in myoelectric control (Phinyomark and Scheme, 2018). By comparing a sampling of conventional machine learning, time series, and deep learning architectures trained to learn the temporal structure of each gesture, this work analyzes the current challenges of zero-shot discrete myoelectric control by answering the following questions:

1. What are the *baseline user-dependent accuracies* for discrete myoelectric control? (Section 3.1)
2. Can *multi-subject cross-user models* generalize to new unseen users? (Section 3.2)
3. How does the *selection of window and increment size* affect system performance? (Section 3.3)
4. How many *subjects and repetitions per subject are needed* for robust zero-shot cross-user models? (Section 3.4)
5. What *features* and/or *feature sets* generalize well to discrete cross-user models? (Section 3.5)
6. Are there bias effects when *end-user demographics are under-represented* in the training set? (Section 3.6)
7. Can model robustness be improved by *limiting the gesture set*? (Section 3.7)
8. Can *transfer learning* approaches be used to fine-tune models to the end-user? (Section 3.8)
9. Do cross-user models *translate to a completely separate dataset* (using the same device)? Additionally, are they *more resilient to common confounding factors*? (Section 3.9)
10. What is the impact of *confidence-bas*ed rejection on within-set and out-of-set inputs? (Section 3.10)

By answering these questions, this work seeks to move the field into a new generation of discrete myoelectric control, validating and analyzing the use of large multi-user models to enable zero-shot EMG-based gesture recognition.

## 2 METHODS

### 2.1 Data Acquisition

This work leverages two separate datasets: (1) the publicly available EMG-EPN612 (N=612) and a novel (2) Myo DisCo (N=14) dataset, both of which were recorded with the Myo Armband, a previously commercially available 8-channel surface-EMG device that samples at 200 Hz (Rawat et al., 2016; Benalcázar et al., 2017). Consistency in electrode location for both datasets was loosely enforced by asking participants to place the device proximal to the elbow, with the charging port facing down on the extensor muscle, to encourage use in *approximately* the same position across users. The default training position for both datasets was the user sitting in a chair with their arm bent at 90 degrees. Participants went through a screen-guided training session where they were shown each gesture and had a certain amount of time allotted for eliciting each gesture (2-5s). All gestures were discrete inputs, meaning the user started at rest, transitioned into an active class, and returned to rest in a relatively short period (*∼*1-2s). In both cases, no feedback was provided to participants. An in-depth description of each dataset follows.

#### 2.1.1 Dataset 1: EMG-EPN612

The EMG-EPN612 (see Figure 1) is a publicly available dataset consisting of 612 participants eliciting the six gestures that were made available by the Myo Armband (Rest, Wave In/Wrist Flexion, Wave Out/Wrist Extension, Hand Open, Hand Close, and Double Tap/Pinch) (Benalcazar et al., 2021). Each dynamic gesture was elicited at some point in a five-second data collection window, with the start and end indices being included as metadata. These indices were used to segment the discrete gestures from the longer five-second recording. The ‘rest’ gesture templates, which were five-second recordings, were cropped before training so that their lengths were generally consistent with those of the other class templates. Also included in this dataset is the native continuous decision stream output by the original Myo device for each gesture. The authors of the dataset have split it into predefined training and testing sets of 306 participants each. Although both sets included 50 repetitions of each gesture, half of the labels were not made available for the testing set as the authors kept them private for an internal competition. Correspondingly, this work omits these data, meaning only 25 repetitions from the testing set were used in the following analyses.

**Figure 1.**
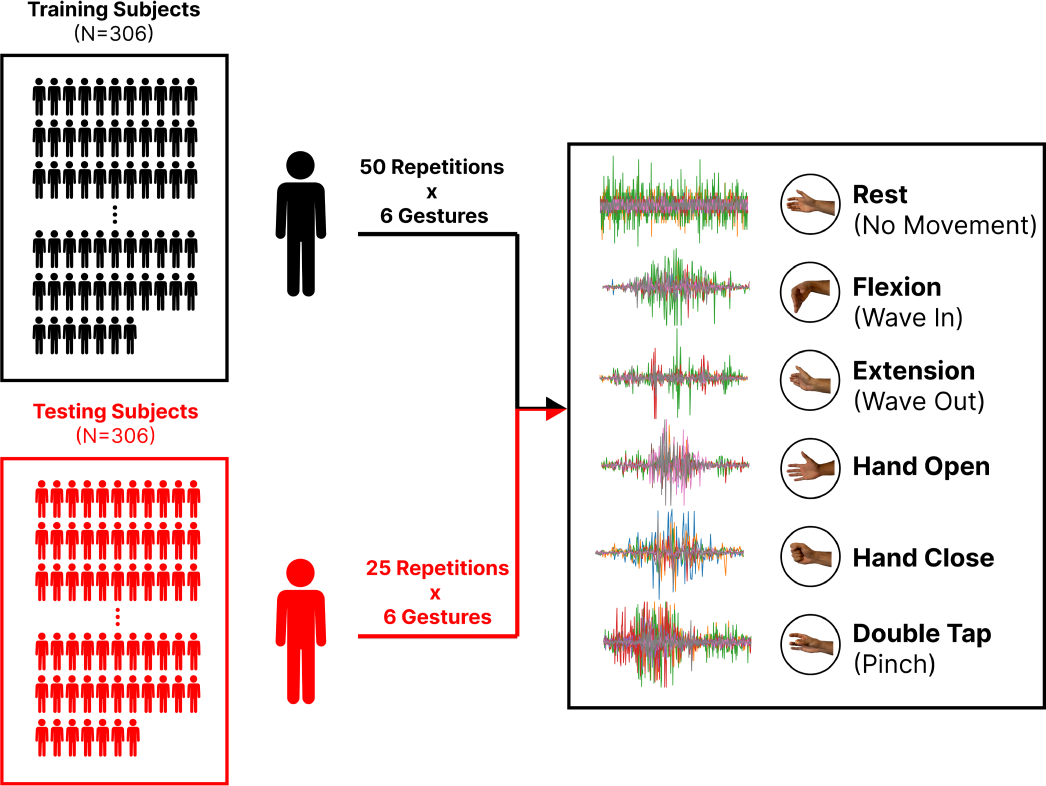
The EMG-EPN612 dataset is split into training (N=306) and testing (N=306) sets for a total of 612 subjects. All subjects in the training group elicited 50 repetitions for each of the six gestures (Rest, Flexion, Extension, Hand Open, Hand Close, and Double Tap). Only 25 repetitions for each gesture are available in the testing set. Also shown is an example of the raw EMG (after being cropped) for each gesture.

#### 2.1.2 Dataset 2: Discrete Confounding Factors (Myo DisCo)

To further evaluate the performance of the large cross-user models developed using the EMG-EPN612 dataset and explore their generalizability to completely unseen data and conditions, a separate dataset — Myo (Dis)crete (Co)nfounding Factors — with intentionally introduced confounds was recorded. Fifteen users participated in this novel collection as approved by the University of New Brunswick’s Research Ethics Board (REB 2023-140). Following a similar protocol to that of the EMG-EPN612 dataset, twenty repetitions of ten gestures: Hand Open, Hand Close, Wrist Flexion, Wrist Extension, Double Tap, Finger Snap, Thumbs Up, Index Point, and Finger Gun were recorded across two sessions occurring between one and seven days apart. Additionally, five repetitions of the gestures were recorded in four additional limb positions (hand by hip, hand above the shoulder, hand in front of the chest, and hand extended away from the body), and ten repetitions of each gesture were recorded at different speeds (50% faster and slower than the user’s comfortable speed). The goal of acquiring these data was to test generalization to a new dataset and to evaluate the effect of three confounding factors: (1) cross-day use, (2) limb position variability, and (3) speed, on cross-user model performance and the impact of out-of-set inputs (i.e., other gestures). Unlike the EMG-EPN612 dataset that was pre-segmented, this dataset was segmented using an automated active thresholding strategy using the “rest” class as a baseline. A further description of this dataset can be found in (Eddy et al., 2024b). Note that one participant of the original N=15 was excluded after it was found that the armband had been oriented in the wrong direction for their collection.

### 2.2 Discrete Control

As defined in (Eddy et al., 2023c) and reinforced by Ctrl Labs (Labs et al., 2024), discrete myoelectric control treats the entire evolution of a dynamic gesture as one input, leading to a single gesture prediction (or event) as the output. Each discrete gesture is thus linked to a single action, similar to a button press. For example, a swipe of the wrist may dismiss a phone call, a double tap of the thumb and index fingers may silence an alarm or directional thumb swipes could be used to guide a character through a virtual obstacle course (e.g. instead of using the arrow keys on a keyboard) (Labs et al., 2024). This is fundamentally different from the continuous control approach that has been commercialized for controlling powered prostheses (Englehart and Hudgins, 2003) whereby the classifier is constantly making predictions at a fixed increment to micro-adjust a prosthesis. In this way, the term *discrete* applies temporally, in that it refers to the generation of specific outputs based on event-driven inputs, in contrast to the more traditional decision stream of *continuous* outputs. In this way, discrete control follows a many-to-one mapping, whereas continuous control follows more of a one-to-one mapping (as in one window of data to one output). This should not be confused with an alternative definition, which refers to the spatial division of a feature space into discrete categories when comparing classification-based with more continuous regression-based approaches (Xiong et al., 2024). In this work, three different algorithmic approaches to achieving this discrete event-based control are explored: (1) Majority Vote LDA (MVLDA), a conventional machine learning approach; (2) Dynamic Time Warping (DTW), a common distance-based time series approach; and (3) Long Short-Term Memory (LSTM), a popular deep-learning time series approach. Additionally, a selection of EMG features — Root Mean Square (RMS), Mean Absolute Value (MAV), Wavelet Energy (WENG), Mean Power (MNP), Waveform Length (WL) — and feature sets — Hudgin’s Time Domain (HTD) and Low Sampling 4 (LS4) — as made available through LibEMG, are explored (Eddy et al., 2023b). Unless otherwise specified, the RMS feature was used as the baseline for each model. These approaches are described in more detail as follows.

#### 2.2.1 The Myo Armband

The Myo Armband was a previously commercially available, low-cost, eight-channel surface-EMG device (Rawat et al., 2016; Benalcázar et al., 2017). Although it has now been discontinued, it was quite influential after its release in 2014 as it lowered the cost and expertise needed to explore EMG-based control (Eddy et al., 2023c). Through proprietary software, the device enabled the recognition of five gestures: Hand Open, Hand Close, Wave In (Flexion), Wave Out (Extension), and Double Tap, which users could calibrate by recording a single repetition of each in a start-up tutorial. The armband also had an SDK with ‘on gesture callbacks’, meaning researchers could develop EMG interfaces without machine learning or signal processing expertise. However, many found that the built-in gesture recognition system did not work well (Kerber et al., 2015; Torres, 2015; Honorof, 2015), leading to negative experiences for the uninitiated and thus poor general impressions of myoelectric control. To the best of our knowledge, this software’s source code has never been made publicly available, meaning that the underlying model remains unknown. Fortunately, the EMG-EPN612 dataset includes the continuous labels from the built-in Myo Armband gesture recognition tuned for every user using one calibration repetition of each gesture. To obtain a discrete output from the continuous decision stream of the Myo, the mode of the active predictions for a given gesture was assumed to be the single discrete label.

#### 2.2.2 Majority Vote Linear Discriminant Analysis (MVLDA)

As described by (Saponas et al., 2008) and (Eddy et al., 2024b), the majority vote approach to discrete gesture recognition employs the legacy continuous classification approach but then converts the decision stream to a single discrete label by computing the mode of the predictions across all windows extracted from the entire gesture template. In this work, a linear discriminant analysis (LDA) classifier was used as the backbone for this majority vote approach, as it is commonly used in prosthesis control as a baseline condition for comparison (Botros et al., 2022; Duan et al., 2021). For all analyses using the MVLDA, the window length from which classifier decisions were made was 200ms (40 samples) with increments between decisions of 25ms (5 samples), as is typical for continuous myoelectric control (Smith et al., 2010). The length of the elicited gesture, as determined by the start and end index metadata (see Section 2.1.1) or by the active thresholding approach (see Section 2.1.2), dictated the length of the gesture which directly influenced the number of windows considered by the majority vote (i.e., longer gestures have more windows).

#### 2.2.3 Dynamic Time Warping (DTW)

Dynamic time warping is a technique used to compare the similarity between two time series while allowing for differences in alignment and length (Sakoe and Chiba, 1978). This is particularly beneficial for discrete gesture recognition, as different repetitions of the gestures will inevitably vary in length, whether inadvertently or intentionally (see Section 3.9). For time series classification (i.e., the classification of a sequence using a single prediction label), the one-nearest neighbour (1NN) DTW classifier has become the baseline for comparison, whereby the training sample with the closest DTW distance to the input is assigned as the classifier’s output (Bagnall et al., 2017). Similar approaches have been previously adopted in EMG-based gesture recognition, such as for hand-writing (Li et al., 2013; Huang et al., 2010) and wake-gesture recognition (Kumar et al., 2021; Eddy et al., 2024a). It has also been recently shown that DTW-based approaches can even sometimes outperform deep-learning approaches when training user-dependent models for recognizing nine discrete gestures (Eddy et al., 2024b). Using the `tslearn` time-series implementation of DTW distance (Tavenard et al., 2020), the 1NN DTW was adopted as a baseline for time series approaches in this work. Based on previous work (Eddy et al., 2024a), the root-mean-square (RMS) feature, extracted from non-overlapping windows of 25ms (5 samples), was used as the input to the DTW-based classifier. These shorter window lengths and increments were selected because the DTW can leverage the time-varying dynamics of the EMG signal to its advantage during prediction.

#### 2.2.4 Long Short-Term Memory (LSTM)

As gesture recognition research continues to evolve, so too does the robustness and complexity of the deep learning models used, meaning that the state-of-the-art is always changing. While other temporal architectures exist and have been shown to have success for EMG-based gesture recognition (e.g., transformers (Zabihi et al., 2023) and convolutional neural networks with reinforcement learning (Vásconez et al., 2023)), the foundational LSTM deep temporal network was adopted in this work to establish the impact of the big data approach, knowing that these results should generalize and may possibly improve with more complex models and architectures. As described in Figure 2, the LSTM model consisted of three LSTM layers of 128 neurons each. After splitting the discrete gesture into non-overlapping windows of 25ms (5 samples) and extracting the RMS feature, as done with the DTW model, the three LSTM layers projected the *N* windowed template to a single 128-dimensional point in the latent space. Note that all gestures were buffered with zeros to the maximum gesture size in the dataset, but the output at the actual end of the gesture was used to compute the loss (Eddy et al., 2024b). After passing through the LSTM layers, the one-dimensional embeddings were passed through three linear layers of 128, 64, and 6 neurons, and finally through a sigmoid function. Instance normalization was applied between these layers. Cross Entropy loss, an Adam optimizer, and a scheduler (*step=5* and *gamma=0.9*) with an initial learning rate of 1e-4 were used to train the model. For the cross-user models, a batch size of 1000 was used during training. Additionally, using a validation set of six participants, early stopping was employed once the validation accuracy did not increase by at least 0.1% for five epochs. For the user-dependent models, a batch size of 200 and a validation set of five repetitions per gesture were used during training. The early stopping strategy was the same, except it only started after 50 training epochs had been completed. All code was developed using PyTorch (Paszke et al., 2019).

**Figure 2.**
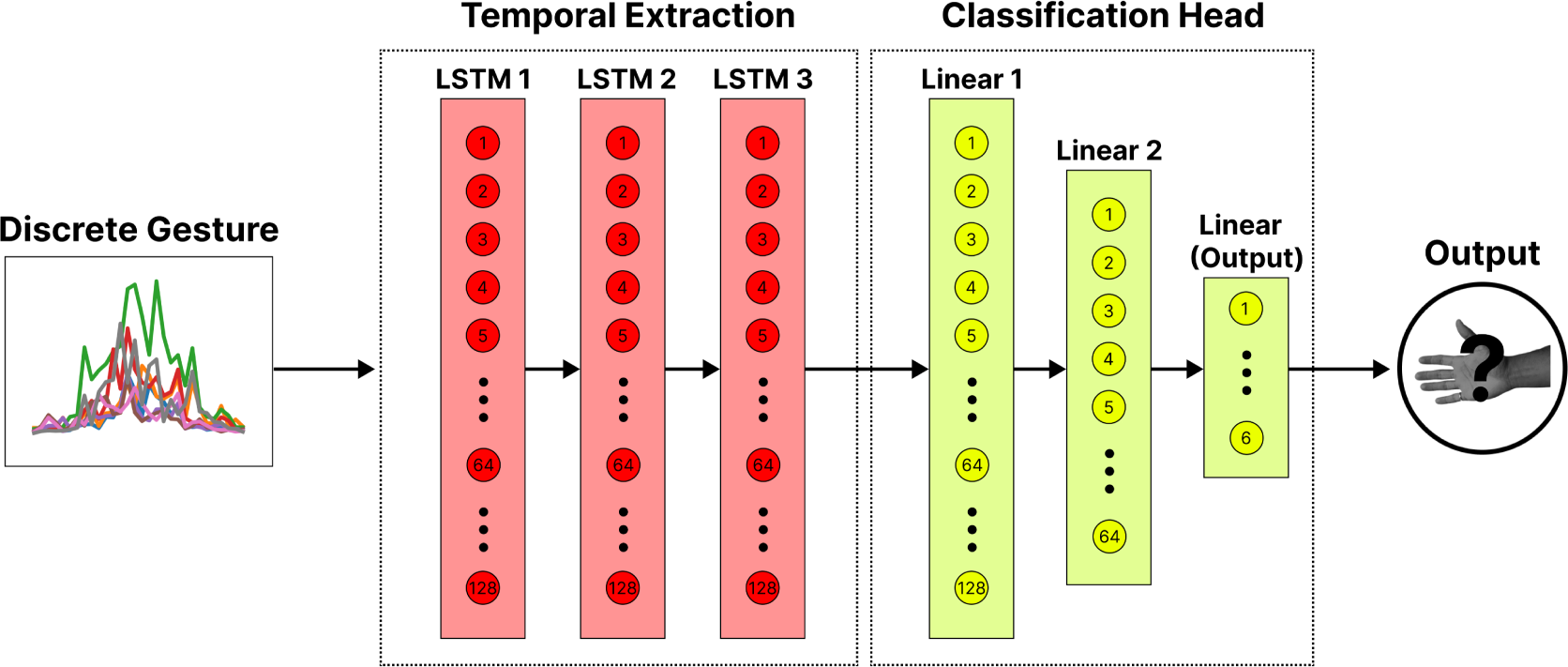
The LSTM-based deep network for classifying discrete gestures. The windowed gestures (with the RMS feature extracted) are passed through three LSTM layers of 128 neurons. This temporal extraction block projects the multi-dimensional gesture template to a single point in the embedding space. The output from this block is then passed through a classification head consisting of two linear layers of 128 and 64 neurons and a final output layer of 6 neurons (corresponding to the number of unique gestures).

## 3 FINDINGS

This section highlights the various findings of this work, answering the questions highlighted in Section through a series of mini-studies. All statistical analyses were conducted using the Statistical Tests for Algorithms Comparison (STAC) online platform (Rodŕıguez-Fdez et al., 2015).

### 3.1 User-Dependent Models

#### 3.1.1 Methods

This analysis was conducted to establish a conventional user-dependent (i.e., within-user) baseline performance for a set of different discrete architectures. A user-specific model was trained and tested for each of the 306 testing users from the EMG-EPN612 dataset following a leave-one-repetition-out cross-validation approach using 20 (of the 25) repetitions. The extra five repetitions of each gesture were saved as a validation set for training the deep learning model. Results are reported as the average classification accuracy (%) across the 20 repetitions of each gesture across all users. Only the 306 testing users were included in this analysis to enable a direct comparison of results with the later cross-user (user-independent) models described in Section 3.2. Section 2.2 provides more information on the training procedure for each model. A repeated measures ANOVA with a Bonferonni-Dunn post-hoc analysis was used to check for statistical significance (*p <* 0.05) between algorithms.

#### 3.1.2 Results

The performance of the user-dependent models is summarized in Figure 3. Each of the tested models significantly outperformed (*p <* 0.05) the built-in gesture recognition decisions output by the Myo Armband (algorithm unknown). While the two temporal models (LSTM and DTW) significantly outperformed the continuous MVLDA approach when using the RMS feature, differences were not significant when using the HTD or LS4 feature sets.

**Figure 3.**
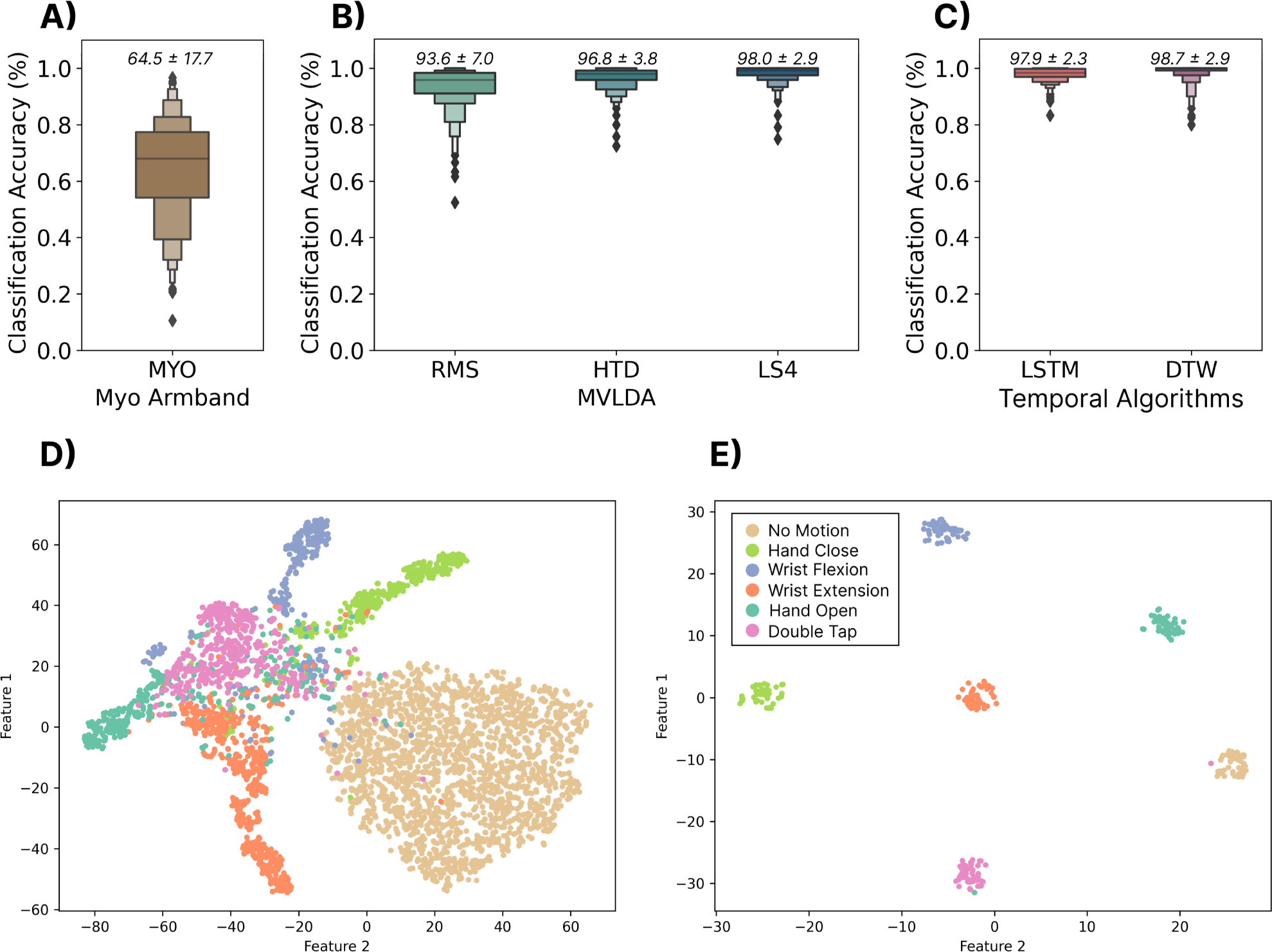
**Top Row:** The user-dependent performance (using the 306 testing subjects) for three commonly used discrete models. The classification accuracy was computed using a leave one-repetition out cross-validation approach on the 20 repetitions of each gesture. **(A)** Shows the performance of the built-in gesture prediction of the Myo that was calibrated with one repetition of each gesture. Above each boxplot is the mean and standard deviation. **(B)** Shows the performance when using a majority vote over the traditional continuous classification approach for three sets of features—root mean square (RMS), Hugins’ time domain feature set (HTD) and low sampling 4 (LS4) feature set. **(C)** Shows the performance of the two sequence modelling approaches—long short-term memory (LSTM) and dynamic time warping (DTW). **Bottom Row: (D)** The t-SNE feature space for a single subject for individual windows of 200 ms with 100 ms increments, computed with the LS4 feature set. **(E)** The t-SNE (t-distributed Stochastic Neighbor Embedding) feature space of the output embedding of the LSTM model.

#### 3.1.3 Discussion

The results from this section show that user-dependent discrete myoelectric control systems can achieve high classification accuracies (*>*97%) when using user-specific offline data. Corroborating previous work (Eddy et al., 2024b), the simple 1-NN DTW classifier outperformed the deep LSTM model, indicating that there may have been insufficient within-user training data to fully leverage the deep-temporal models. Both temporal models (i.e., DTW and LSTM) that used the RMS of the signal as input yielded significantly higher accuracies than the majority vote approach when using the same feature. When more descriptive feature sets were used (e.g., HTD and LS4), the performance was much more comparable, suggesting that there were enough steady-state portions in these discrete gestures for majority-vote approaches to work. However, in the case of more dynamic gestures, such as sign-language words or handwriting recognition (Labs et al., 2024), these approaches may fail. The performance of the built-in gesture recognition of the Myo Armband (the model of which is unknown) was comparatively quite poor, significantly under-performing all other approaches. Therefore, it makes sense that many individuals, particularly those who had previously used the default Myo control, believed the technology lacked sufficient robustness for real-world use (Karolus et al., 2022; Eddy et al., 2023c). It should be noted that while otherwise high accuracies were achieved, these results were obtained using a relatively constrained testing set (i.e., the user sitting in a chair with all gestures elicited in the same limb position). This is unlikely to represent real-world use cases where confounding covariate factors, like cross-day use, limb position variability, and gesture elicitation speed, will inevitably be present (see Section 3.9). Moreover, requiring users to undergo 12.5 minutes of data collection (25 reps x 6 gestures x 5 seconds) every time the armband is donned to achieve such performance is an unrealistic expectation. Consequently, although these results serve as a promising user-dependent baseline, the rest of the paper focuses on enabling robust cross-user discrete gesture recognition whereby no data is required from the end user.

### 3.2 Many-Subject Cross-User Models

#### 3.2.1 Methods

In this section, the viability of zero-shot discrete gesture recognition using large cross-user models was explored. To do this, all 50 repetitions from each of the 306 training users from the EMG-EPN612 dataset were used to train a single cross-user model. For the deep LSTM model, the training users were further split into a 300-user training set and a 6-user validation set. As before, all models were then evaluated using all of the data from the 306 EMG-EPN612 testing users. This case represents the zero-shot, cross-user performance, meaning that the models had not previously seen any data from the test users, simulating the ideal ‘plug-and-play’ scenario. Although the same MVLDA, DTW, and LSTM approaches were evaluated as in the user-dependent case, a limitation of DTW-based architectures is that their complexity scales linearly with the number of templates in the training set. In this case, there were 306 training subjects and 50 repetitions of each gesture, meaning there were 91,800 (306 users x 50 reps x 6 gestures) training templates in the dataset. Using a standard nearest-neighbour classification approach, each new template would have to be compared to all 91,800 templates, which is computationally prohibitive for real-time control. Correspondingly, two alternative solutions were explored to make cross-user DTW computationally feasible. For the DTW (Closest) case, the closest template to all other templates in the dataset (the one whose sum of DTW distances to all other templates was minimum) was selected as a single representative template for each gesture. For the DTW (Mean) case, all templates for each gesture were interpolated to be the same size, and then the mean template was computed. This meant that for both cases, there were only six target templates against which to compare (1 for each possible class). The MVLDA and LSTM models were trained as described in Section 2.2 without algorithmic changes other than the appropriate selection of training data. A repeated measures ANOVA with a Bonferonni-Dunn post-hoc analysis was used to check for statistical significance (*p <* 0.05) between algorithms.

#### 3.2.2 Results

Figure 4 summarizes the cross-user performance for all models. The LSTM significantly (*p <* 0.05) outperformed the MVLDA approaches by 23.9%, 18.4%, and 17.4% when using the RMS, HTD, and LS4 features. Additionally, although the DTW performed well for the user-dependent case, the modified strategies did not scale well for cross-user models. In this case, the LSTM significantly outperformed (*p <* 0.05) the DTW (Closest) and DTW (Mean) approaches by 18.2% and 18.9%, respectively. The effect of the feature sets on the performance of the MVLDA models was similar to the user-dependent case (6.5% difference for the LS4 set here, compared to the 4.4% difference in the user-dependent case). Finally, Figure 4D shows a t-SNE projection of the distributions of the features extracted from the windowed data. The high degree of overlap between gestures in this ‘continuous’ feature space may explain the mediocre performance of the majority vote approaches when scaled to cross-user models. In contrast, the t-SNE projection of the embedding space from the LSTM model (Figure 4E) shows much better separability.

**Figure 4.**
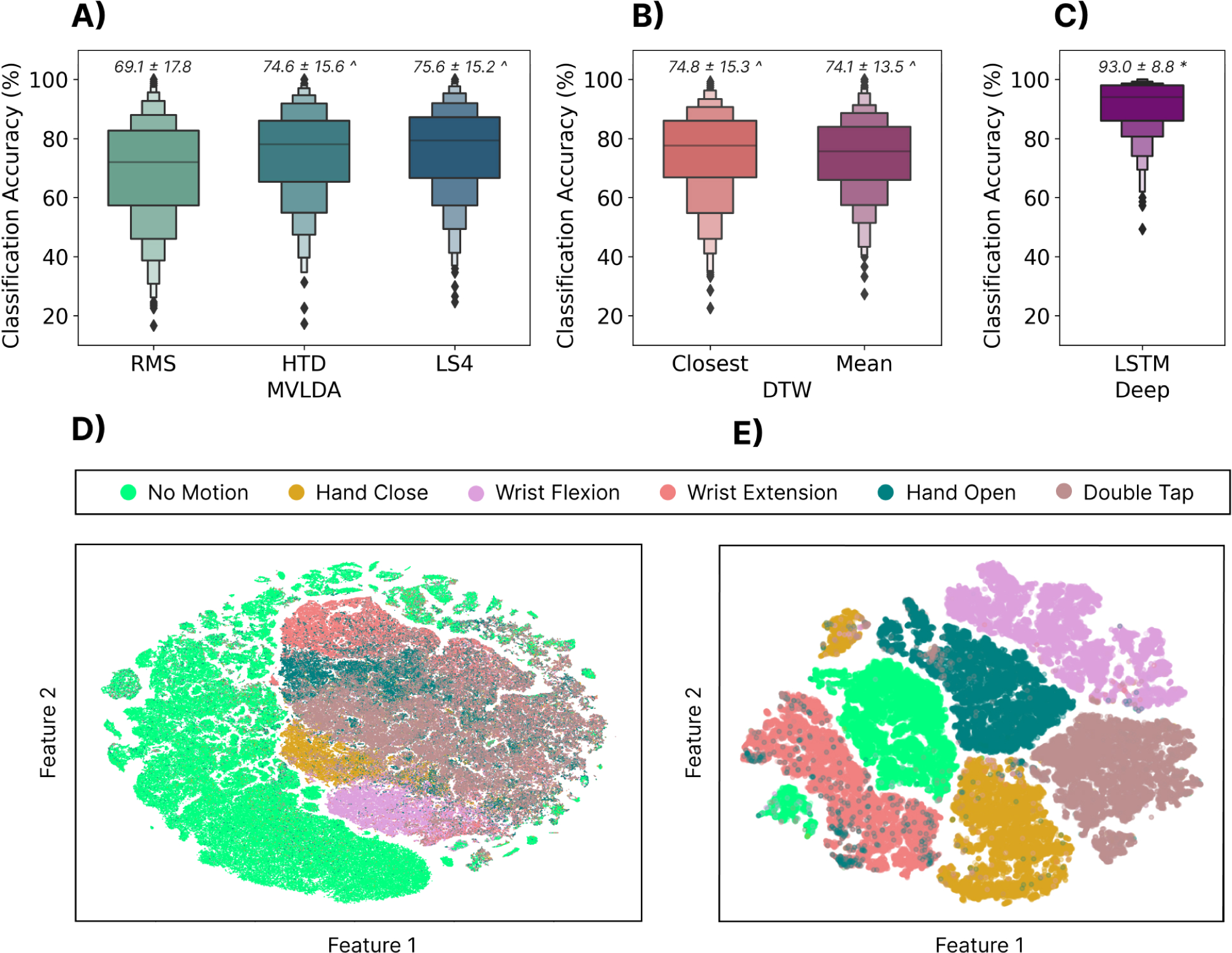
**Top Row:** The zero-shot cross-user classification accuracy when training with the 300 training subjects and testing on the 306 testing subjects for three different types of models. **(A)** The cross-user MVLDA model trained on the RMS, HTD, and LS4 features. **(B)** The DTW (closest) model used the template for each gesture with the minimum distance to all other templates. The DTW (Mean) model used the mean gesture templates across users as the target template. **(C)** The cross-user LSTM model. Above each boxplot is the mean and standard deviation where * and *∧* indicate significant differences (*p <* 0.05) to all other algorithms and MVLDA (RMS), respectively. **Bottom Row: (D)** The t-SNE-projected feature space for individual windows of the RMS feature. **(E)** The t-SNE-projected feature space of the output embedding of the cross-user LSTM model.

#### 3.2.3 Discussion

The standard for discrete myoelectric control was arguably established in 2008 by Saponas et al. when they showed the majority vote approach applied to a continuous decision stream for recognizing discrete finger taps (Saponas et al., 2008). Although this approach and the idea of *muscle-computer interfaces* seemed promising at the time, classification performance was shown to significantly deteriorate (*∼*57%) when testing on unseen users (Saponas et al., 2008). Even with many more subjects (N=300), the results from this work corroborate that majority vote-based approaches based on the continuous decision stream of a stationary classifier do not scale well for cross-user models.

Although DTW-based approaches worked well for user-dependent models, the results from this work suggest that they also do not scale well for cross-user models, albeit with a few caveats. This work applied two simple attempts to generalize DTW to the cross-user case, and performance improvements are likely still possible. In particular, given the performance of DTW in Section 3.1, exploring template blending approaches or multi-template implementation improvements may lead to better cross-user DTW-based models, and is an interesting area for future research. Nevertheless, these results suggest there is enough variability between user templates (whether for physiological or behavioural reasons) that they cannot effectively be described by a single representative template.

The deep-LSTM model significantly outperformed the other approaches, with classification accuracies of 93.0%. This performance improvement is likely driven by two factors: (1) deep learning models benefit from more data, such as in the case of the EMG-EPN612 dataset, and (2) LSTMs inherently model the temporal evolution of each gesture, meaning that its onset, offset, and everything in-between, help define the embedding space. Compared to the majority vote approach based on the continuous decision stream, the LSTM learns each gesture’s temporal structure, which is likely more similar across participants (compared to individual out-of-context windows), thus leading to improved performance. Additionally, unlike DTW, neural networks are capable of modelling multi-modal distributions without a linear increase in computation time, enabling them to learn and model behavioural and physiological differences in user data while maintaining quick inference times for real-time control.

### 3.3 Effect of Window and Increment Length

#### 3.3.1 Methods

This analysis aimed to understand the impact of different window lengths and increments (i.e., the temporal resolution of the gesture template) on cross-user classification performance. The classification accuracies for combinations of a set of window lengths between 5 and 77 samples (25 ms and 385 ms) and increment sizes between 1 and 77 samples (5 ms and 385 ms) were calculated. The 77-sample upper limit was selected based on the shortest gesture template in the dataset. For all combinations of window lengths and increments, the maximum number of windows that could be fit using those parameters were extracted from each gesture template. For example, a gesture template that was 100 samples long would yield 3 windows when the window length and increment are 77 and 10 samples, respectively. The training and testing sets were split as highlighted in Section 3.2. A depiction of the temporal profiles (i.e., the RMS values extracted from a sequence of windows) of a representative discrete gesture (hand open) is shown in Figure 5 for a selection of different window and increment lengths.

**Figure 5.**
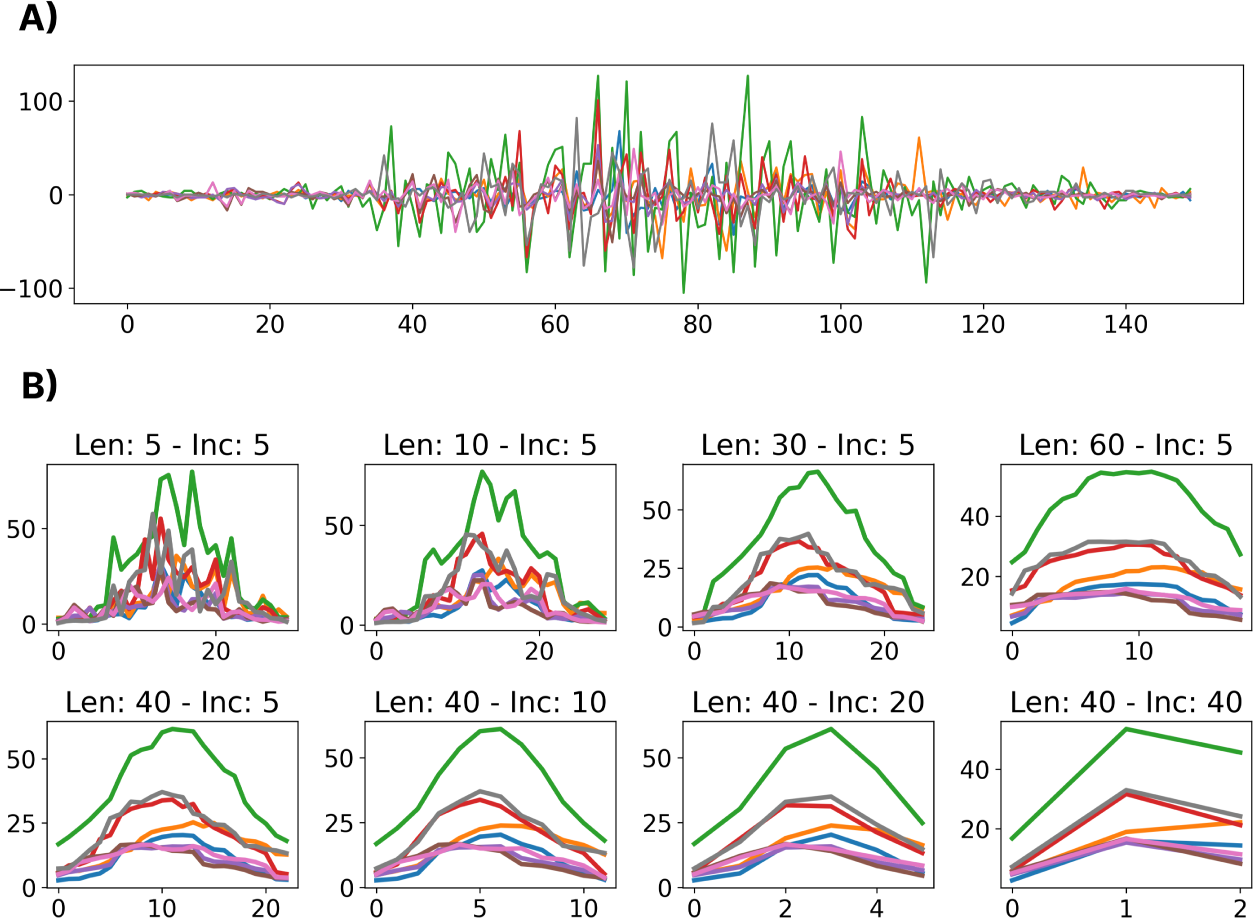
A visualization of the effects of different window lengths and increments applied to the same gesture. **(A)** Shows the raw EMG for a representative dynamic gesture from the dataset. **(B)** Shows the extracted RMS of the windowed gesture for a variety of window lengths (Len) and increments (Inc).

#### 3.3.2 Results

As shown in Figure 6A, the MVLDA approach benefits from increased window lengths as they improve the RMS estimate (the signal is assumed to be stationary within each window). This is seen in the lower portion of the heatmap (larger window lengths), where the highest classification accuracies were obtained.

**Figure 6.**
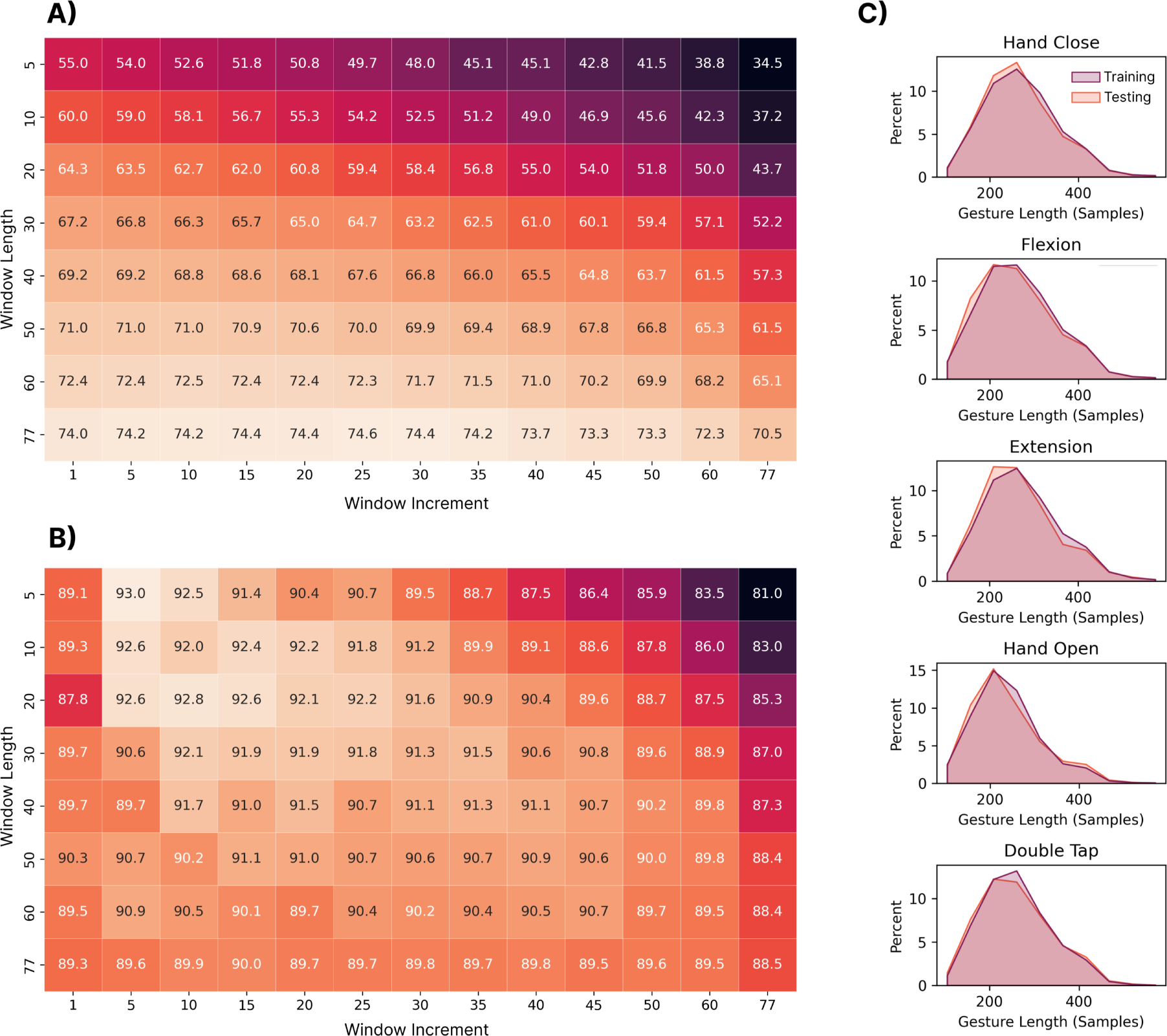
(**A**) The impact of the window length and increment on the performance of the **MVLDA** with the RMS feature and **(B) LSTM** discrete approaches. Each box corresponds to the classification accuracy when trained on the 306 training users and tested on the 306 testing users. The window length and increment values correspond to the number of EMG samples (1 sample at 200 Hz = 5 ms). **(C)** Shows the distribution of gesture lengths (i.e., number of samples) of each gesture across all training and testing subjects.

It is worth noting that the combination of longer window lengths and increments begins to degrade performance. This is because there begins to be very few windows from which to compute the majority vote and, for the shortest gestures, only a single window (from the early portion of the sequence) may be extracted. In contrast, the deep LSTM model (see Figure 6B) benefited from shorter windows and increments as this enabled it to better model the temporal dynamics of each gesture. Correspondingly, the upper left-hand side of the heatmap (small window lengths and increments) shows the highest classification accuracies. The best window and increment size combinations were: (1) window length = 77 samples and window increment = 25 samples for the MVLDA (74.6%) and (2) window length = 5 samples and window increment = 5 samples (93.0%) for the LSTM. A Shapiro-Wilk test for normality (*W* = 0.99, *p <* 0.0001) showed that the length of the gestures (i.e., the speed with which they are elicited) in the training and testing sets are roughly equal and normally distributed (see Figure 6C), with the average gesture length being 1.3 seconds.

#### 3.3.3 Discussion

Although the impact of window length and increment have been widely explored for continuous myoelectric control (Smith et al., 2010), their role in predicting temporally-driven discrete EMG-based gestures has received less attention. Generally, the results found here suggest that the stationary models, which do not model the temporal evolution of the gesture, benefit from longer windows from which more smoothed estimates of the RMS could be extracted. In contrast, the LSTM model benefited from shorter windows and increments, providing less smoothed estimates of RMS and higher resolution in the sequence of features that it models. This is an important reminder that care should be taken when making design decisions in discrete control based on knowledge from the broader continuous myoelectric control literature. Future work should evaluate the impact of window length and increment selection when using devices with higher sampling rates that can conceivably provide higher temporal resolution (the Myo device samples at only 200Hz, whereas EMG is typically known to contain frequency information up to 500Hz (Eddy et al., 2023c)).

A secondary outcome of this analysis was the evaluation of the average gesture lengths across participants. Intuitively, the speed at which gestures were elicited (no temporal guidance or feedback was provided) followed a normal distribution, with an average length of 1.3 seconds. Ultimately, this has implications for the design of interactive systems that may use the models trained using this dataset. For example, a scrolling application may have a relatively low bandwidth if the system can only enable one input every second. If the user wishes to scroll faster, this could be problematic since classification performance degrades when gestures are elicited at faster speeds (Eddy et al., 2024b). Another consideration is those users who fall on either end of the normal distribution, as their performance may be more susceptible to generalization errors (see Section 3.6) using such cross-user models (e.g., individuals with less mobility or hand dexterity) (Eddy et al., 2024b). Future work should explore how to become resilient to discrete gestures elicited at different speeds, either through data aggregation in the original dataset, transfer learning, or data augmentations.

### 3.4 Effect of Number of Subjects and Repetitions

#### 3.4.1 Methods

This analysis explored the impact of the number of subjects in the training set and the number of repetitions per subject. Based on the previous results (see Section 3.2 and 3.3), this and subsequent sections perform the zero-shot cross-user classification using only the LSTM model. Two tests were explored: (1) when training with permutations of 1-300 subjects and testing on 306 subjects (N=300) and (2) when training with permutations of 1-500 subjects and testing on 100 subjects (N=500). For both cases, the subjects used for training were randomly chosen from the pool of available subjects. Due to the variability introduced by this randomization, this process was repeated 10 times for each case (e.g., for the 100-subject case, 10 random groups of 100 subjects were used for training) and the results are reported as the average accuracy across the 10 folds (e.g. each box in the heat map shown in Figure 7). Once the 10 random groups of subjects were chosen, they were held constant while evaluating the impact of the number of gestures included. The testing cohorts for each of the tests remained constant across all cases and folds. Note that for the N=500 test, the maximum number of repetitions used was 25 due to the reduced number of repetitions provided for the test subjects in the dataset.

**Figure 7.**
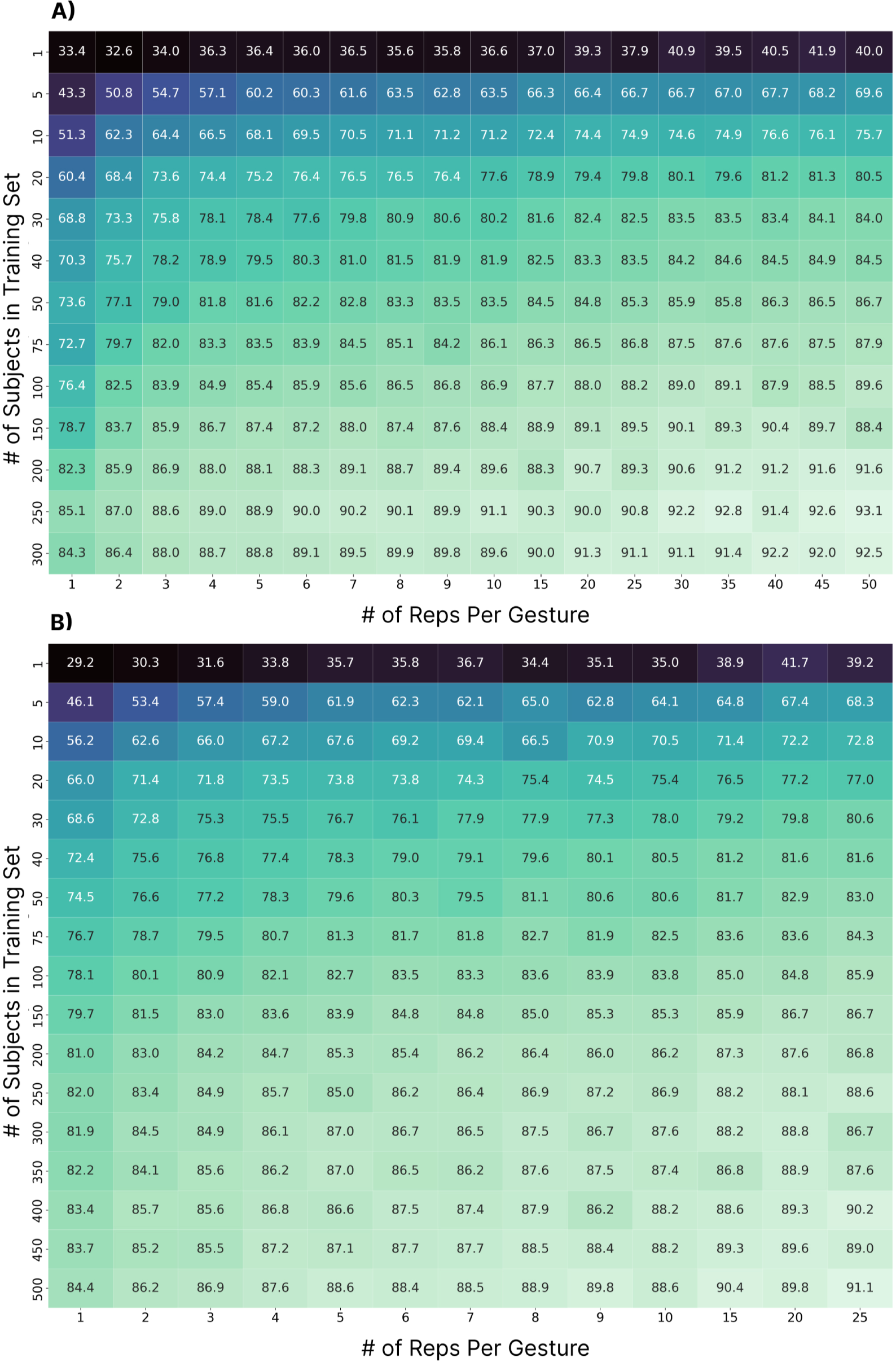
The impact of the number of subjects in the training set and the number of repetitions per subject included for each gesture. Each box shows the average accuracy across ten folds of a random subset of training subjects. **(A)** Represents the case when drawing from the 300 training subjects and testing on the 306 testing subjects. **(B)** Represents the case when using up to 500 subjects for training (drawn from a combination of the preassigned training and testing sets) and 100 subjects for testing.

#### 3.4.2 Results

For the N=300 case (Figure 7A), the cross-user accuracy first surpassed 90% at 150 subjects (with 30 repetitions per gesture), a much higher subject count than what is typically collected for EMG-based gesture recognition research. From a data perspective, this corresponds to, at minimum, 37.5 hours (150 subjects x 30 reps x 6 gestures x 5 seconds) of active data recording, or 15 minutes per user. The accuracy then surpassed the 92% threshold at 250 subjects with 30 repetitions (i.e., 67.5 hours), nearly double the amount of data for a 2% improvement. Even so, the results suggest that additional subjects and repetitions may continue to improve performance, potentially beyond the highest achieved accuracy of *∼*93%. For the N=500 case (Figure 7B), the accuracy achieved the 90% threshold at 400 subjects with 25 repetitions (i.e., 83.3 hours of data). This differed slightly from the N=300 case, but this may be due to small differences in the size of the pool of test subjects from which to draw, or perhaps more importantly, because the testing set contained a disproportionate number of participants (over half) from underrepresented groups in the training set (see Section 3.6). Nevertheless, while the maximum accuracy achieved was 91.1%, the accuracy does not appear to have plateaued, indicating that more subjects may continue to improve performance.

#### 3.4.3 Discussion

Myoelectric control research has historically adopted relatively low numbers of participants due to the experimental burden associated with recruiting and collecting data from many subjects. For example, in Saponas’ early work, one of the first to establish cross-user models as a challenge in HCI, the data from 13 participants and 50 repetitions were used for testing the possibility of zero-shot models (Saponas et al., 2008). Even if all these subjects were used for training, using similar numbers in this work would have yielded a cross-user performance of 75.7% (see Figure 7A). Even with more recent neural networks, when leveraging data from 11 subjects, cross-user continuous myoelectric control performance was found to be only 68%, compared to 92.3% when using transfer learning approaches (Xu et al., 2024). Similarly, in the work by Campbell et al., it was shown that when leveraging ten subjects and 16 reps of ten gestures, naive cross-user models achieved accuracies below 50% (Campbell et al., 2021). These three examples highlight that it has to date been unfeasible to achieve true zero-shot cross-user models with low subject counts. In this work, cross-user models above 90% accuracy were only made possible once 150 subjects and 30 repetitions per gesture were used for training. Recent work by Ctrl Labs suggests that there may be benefit to including as many as 4800 users when training zero-shot cross-user discrete models for recognizing thumb swipes (*∼*7% error rate) (Labs et al., 2024). This is drastically more subjects than current standardized and openly-available datasets such as the 43-participant multi-day wrist/forearm dataset recently released by Pradhan et al. (Pradhan et al., 2022) and the commonly used Nina Pro DB1 and DB2 datasets consisting of 27 and 40 subjects, respectively (Atzori et al., 2015, 2014). Furthermore, a lack of standardized hardware and data collection practices has thus far precluded effective pooling of data across studies. Generally, these results suggest that the challenge in enabling robust cross-user EMG has not been algorithmic alone, but that there has been a general lack of data, supporting the collection and release of large datasets as a crucial future research direction (Phinyomark and Scheme, 2018).

Due to the challenges of collecting large datasets, researchers have turned toward few-shot and transfer learning approaches whereby the backbone of the model leverages other users’ data, and the end model is fine-tuned to the end user (Jiang et al., 2024; Xu et al., 2024; Côté-Allard et al., 2019). While these approaches can reduce some of the training burden associated with myoelectric control, they are not necessarily ideal for the widespread adoption of EMG. For example, it is unlikely that consumers will be inclined to provide training gestures every time they don their EMG device, when competing technologies do not require it. Correspondingly, for the future success of this technology outside of prosthesis control, the EMG community should further explore zero-shot models that work without any training data from the end user (i.e., plug-and-play) or that only require a single, one-time calibration session (not unlike touch or face ID on smartphones).

### 3.5 Feature Selection

#### 3.5.1 Methods

Developing and evaluating handcrafted and deep-learned features has been studied exhaustively in continuous myoelectric control (Phinyomark et al., 2009, 2012; Hudgins et al., 1993; Samuel et al., 2018). Generally, however, these features and feature sets have been optimized for the largely static contractions commonly used to control powered prostheses. Offline studies using user-dependent models have been able to make marginal improvements by introducing increasingly complex feature sets (Khushaba et al., 2022). However, there has been comparatively little exploration of feature selection for cross-user models, and none in the context of discrete myoelectric control. This section evaluates the performance of five common features and two feature sets to better understand the impact of feature selection on discrete cross-user myoelectric control, as demonstrated using the LSTM model. The training and testing sets were split as highlighted in Section 3.2.1. The HTD feature set (Hudgins et al., 1993) was selected as it is often used as a baseline for myoelectric control research, and the LS4 feature set (Phinyomark et al., 2018) as it was designed specifically for low sampling rate devices (such as the Myo). The Friedman aligned-ranks statistical test with Holm correction was used to check for statistical differences (*p <* 0.05) between features and feature sets.

#### 3.5.2 Results

The classification accuracies for all features and feature sets are shown in Figure 8. All had similar performance, achieving accuracies within 1% of each other (between 92.2% and 93.1%). Although some differences were significant, their effect sizes, as measured by Cohen’s D, were all small (*d <* 0.2); RMS-MNP (*d* = 0.09), RMS-MAV (*d* = 0.08), HTD-MNP (*d* = 0.10), HTD-MAV (*d* = 0.09), and WENG-MNP (*d* = 0.09).

**Figure 8.**
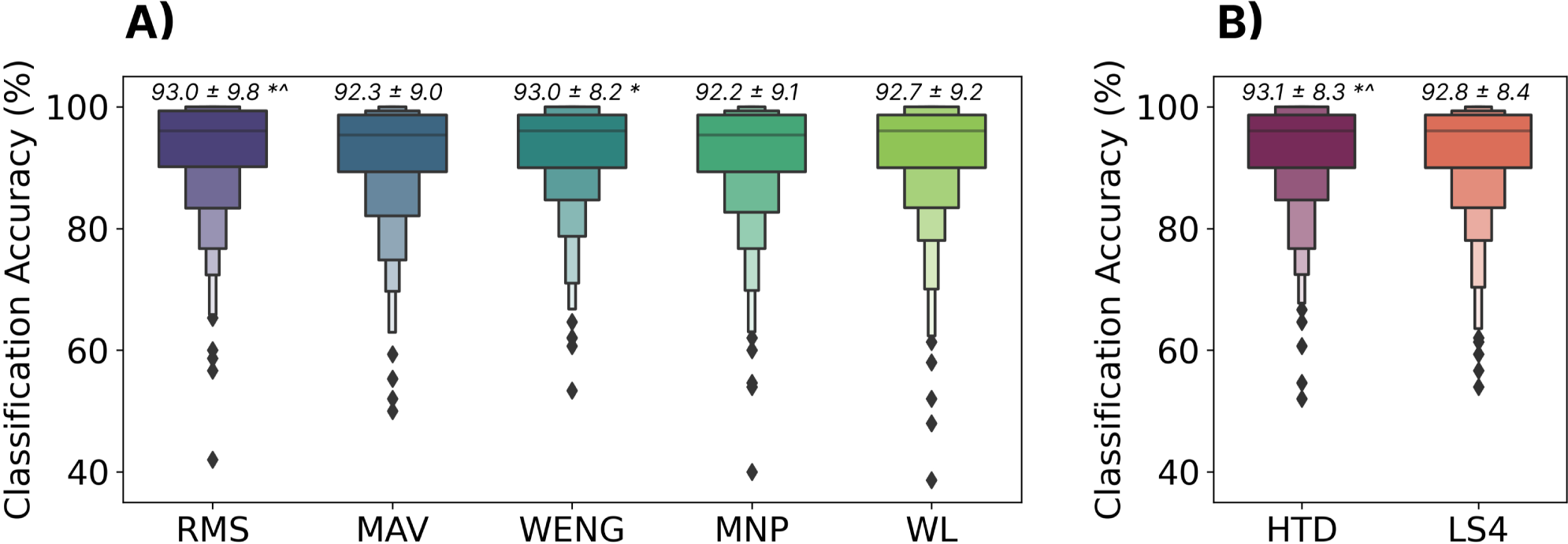
The effect of feature set on zero-shot cross-user performance using the LSTM model. **(A)** Shows individual features: Root Mean Square (RMS), Mean Absolute Value (MAV), Wavelet Energy (WENG), Mean Power (MNP), and WL (Waveform Length). **(B)** Shows two common feature sets: Hudgins’ Time Domain Features (HTD) and Low Sampling 4 (LS4). All features and feature sets are available through LibEMG (Eddy et al., 2023b). Above each boxplot is the mean and standard deviation where * and *∧* indicate significant differences (*p <* 0.05) to the MNP and MAV features.

#### 3.5.3 Discussion

Only small differences (*<*1%) were found between all features and feature sets. This lack of difference, when combined with the superiority of the temporal models as discussed in Section 3.2, suggests that the temporal profile of the gesture is driving the majority of the cross-user classification performance. With the inclusion of the temporal dynamics and small inter-subject differences in behaviours and physiology, it seems that the highly descriptive features usually used to describe differences between individual static windows of EMG (within which the signal is assumed to be stationary) provide little benefit and, in some cases, can even hurt performance. Based on these results, the continued engineering of complex handcrafted features may not be required for discrete myoelectric control and, instead, focus may more appropriately placed on deep approaches for modeling the profile (e.g. transformers (Vaswani et al., 2017) or distortion loss with shape and time (Le Guen and Thome, 2019)).

### 3.6 Effect of Bias in the Training Data

#### 3.6.1 Methods

As previously highlighted, cross-user myoelectric control models have been difficult to achieve partially due to physiological and behavioural differences between individuals. While this work shows that cross-user models are possible when including enough participants, the impact of these differences on model training is still not fully understood. In particular, it was hypothesized that imbalances in the sources of such differences in the training set might lead to biased systems (i.e., systems that favour one physiology or set of behaviours) (Leavy, 2018; Mehrabi et al., 2021). Using the metadata provided by the EMG-EPN612 dataset, this section evaluates the role of two physiological differences on system bias: (1) participant gender (man/woman as reported in the dataset) and (2) handedness (left/right). Additionally, a behavioural difference, gesture elicitation speed, is also evaluated. Such factors have seldom been evaluated in myoelectric control, as most research has explored user-dependent cases where these issues are absent. Through these analyses, this section provides additional insights into the importance of acquiring representative training data for ensuring equitable myoelectric control systems that perform well for all users.

The same zero-shot cross-user LSTM model, trained with the 306 training subjects as described in Section 3.2.1, was used to evaluate these effects. The testing set, however, was split based on each of the evaluated categories. Note that the distribution of gender and handedness was found to be consistent across the testing and training sets. The average gesture length for each participant (excluding no movement) was also computed and split into five groups (empirically, based on a histogram). Unpaired t-tests were used to test for significant differences based on gender and handedness. A Friedman test with Holm correction was used to test for significant differences between gesture lengths.

#### 3.6.2 Results

The results suggest that the proportions of gender (man/woman) and handedness (right/left) were 66%-34% and 96%-4% splits, respectively, highlighting a strong bias toward men and right-handed individuals (see Figure 9). These ratios were consistent across the predefined training and testing sets established by the EMG-EPN612 authors. As exemplified by the significantly higher (*p <* 0.05) accuracies for men and right-handed users, this bias played a significant factor in defining the system’s performance for each group. It is interesting to note that the difference in performance based on handedness was substantially larger, as was the imbalance in the training set.

**Figure 9.**
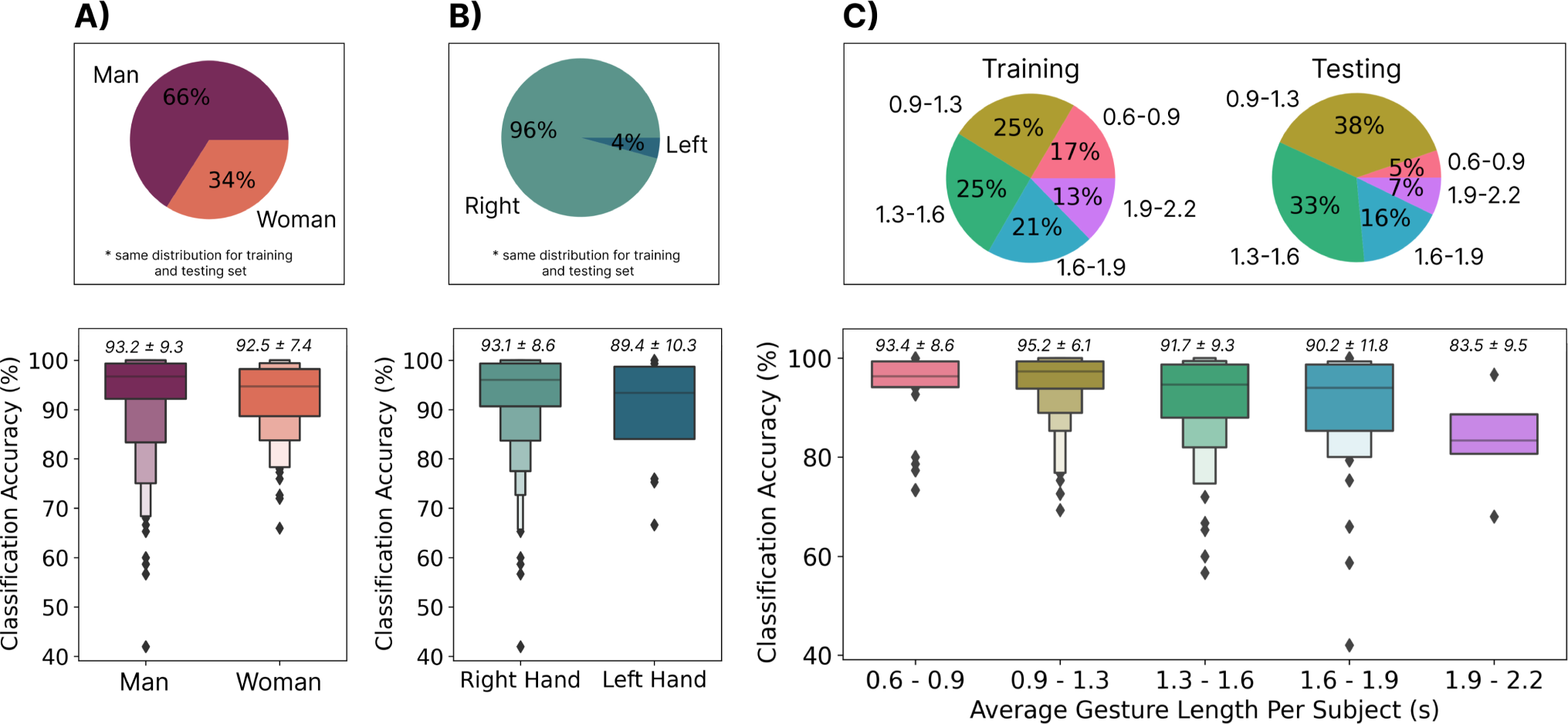
**(A)** The distribution of men and women in the training and testing sets **(top)** and the accuracy of the cross-user model when evaluating each category from the testing set independently **(bottom)**. **(B)** The distribution of right and left-handed users in the training and testing sets **(top)** and the accuracy of the cross-user model when evaluating each category from the testing set independently **(bottom). (C)** The distribution of average gesture lengths per user in the training and testing sets **(top)** and the accuracy of the cross-user model when evaluating each category from the testing set independently **(bottom).**

As highlighted in Section 3.3, the length of gestures followed a normal distribution, with the average length being 1.3s. It thus makes sense that the majority of the distribution for the average gesture lengths for participants fell into bins one (0.9s-1.3s), two (1.3s-1.6s), and three (1.6s-1.9s). The distribution of gesture lengths, as shown in Figure 9, however, shows an unequal distribution of these bins across training and testing. For example, there were fewer shorter gestures (0.6s-0.9s) in the testing set than in the training set. Although significant differences were not found between bins, large differences were found between gesture lengths (up to 11.7% when comparing gestures between 0.9s-1.3s and those *>*1.9s).

#### 3.6.3 Discussion

The EMG-EPN612 dataset did not have an equal representation of gender or handedness, which led to significantly worse performance (*p <* 0.05) when the model was tested on women and left-handed users. First, this suggests that there may be physiological differences in the signal patterns elicited by these groups that should be explored further in subsequent research. More importantly, however, this poses real concerns, including ethical ones, as underrepresented user groups would perform significantly worse when using the system. Similarly, the model performed worse for slower gestures, which could also be a general accessibility issue. For example, individuals with limited mobility (who may benefit most from novel interaction modalities) may take longer than the average of 1.3s to elicit a gesture. It is, therefore, crucial when collecting big datasets in the future that there is an appropriate representation of physiological differences (such as gender and handedness) and behavioural differences (such as gesture elicitation speed). This becomes particularly important when using data from more participants (e.g. the 6400 collected by Ctrl Labs (Labs et al., 2024)), as these biases (such as the gender bias (Leavy, 2018)) could be exacerbated if there remains bias within the training set. Correspondingly, this area deserves increased research attention, and many other physical (e.g., age, ethnicity, and mobility issues) and behavioural factors (e.g., contraction intensity and tempo) should be explored in the context of cross-user EMG models. Combining data from all types of potential users (as further discussed in Section 4) may be crucial for robust and equitable cross-user EMG models; however, the extent to which is currently unknown.

### 3.7 Effect of Gesture Selection

#### 3.7.1 Methods

Although much of the recent literature has endeavoured to expand the number of classes or degrees of freedom that can be predicted by myoelectric control (Atzori et al., 2015; Geng et al., 2016), this increased difficulty ultimately increases the potential for mistakes, working against fears about the robustness of EMG. Conversely, many interactions in HCI could be enabled by a relatively small, but appropriately designed gesture set. For example, scrolling in mixed reality might require three (scroll up, scroll down, and selection) whereas answering or dismissing a phone call might require only two. One compelling opportunity for exploration is thus the selection of gesture set to achieve highly resilient and robust classifiers for such applications in HCI. This section correspondingly explores the zero-shot cross-user LSTM model performance when adopting a variety of gesture subsets. All techniques for model training remained the same for all combinations (see Section 3.2), except that the unused gestures were simply omitted from the training and testing sets. A repeated measures ANOVA with a Bonferonni-Dunn post-hoc analysis was used to check for statistical significance (p*<*0.05) between algorithms.

#### 3.7.2 Results

The classification accuracies for all notable subsets of gestures are shown in Figure 10. Intuitively, reducing the number of gestures tended to improve classification accuracy. In particular, gesture sets 2 (Rest, Open, Close), 3 (Rest, Flexion, and Extension), and 5 (Rest, Flexion, Extension, and Double Tap) yielded significantly better performance (p*<*0.05) than when using all gestures, achieving zero-shot accuracies of *>*96.5%, and as high as 98.2%.

**Figure 10.**
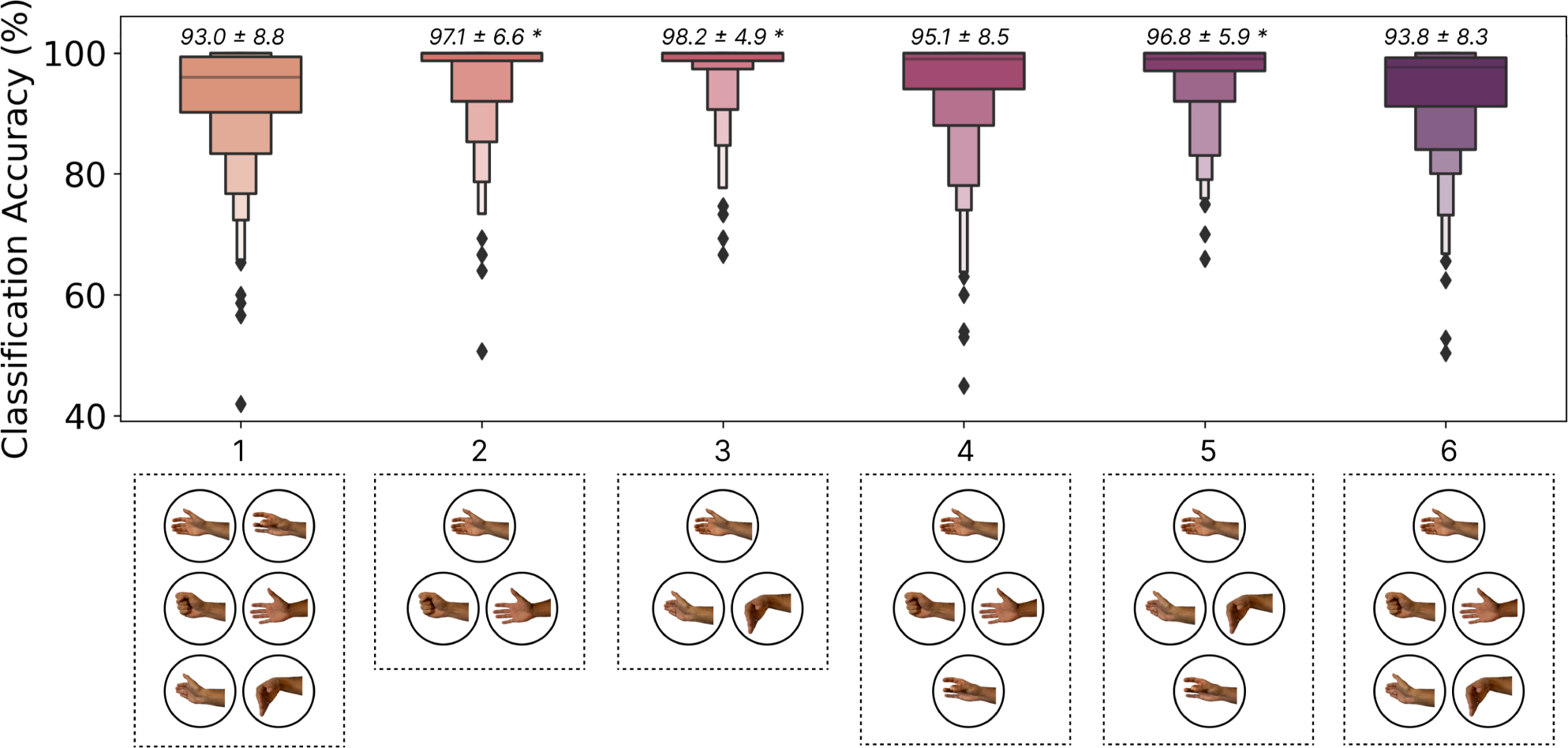
The effect of gesture set selection on the zero-shot cross-user LSTM model performance. The gesture subsets shown, from left to right, were: **(1)** Rest, Flexion, Extension, Open, Close, Double Tap, **(2)** Rest, Open, Close, **(3)** Rest, Flexion, Extension, **(4)** Rest, Open, Close, Double Tap, **(5)** Rest, Flexion, Extension, Double Tap, and **(6)** Rest, Open, Close, Flexion, Extension. Above each boxplot is the mean and standard deviation where * indicates significant differences (*p <* 0.05) with gesture set 1.

#### 3.7.3 Discussion

As exemplified by the commercially available ‘double tap’ gesture on the Apple Watch^1^, emerging HCI applications may benefit from fewer but very robust input gestures. These interactions can then be designed appropriately around the limited input space (e.g., depending on the context, a double tap could scroll through a menu, answer a phone call, or make a selection). The results from this section show that increased accuracy (up to 5.2%) can be achieved by limiting the input space. One possibility is thus to run models trained on a subset of gestures for certain interactions. For example, when a phone call is detected the model trained with gesture set 3 (flexion (answer), and extension (decline)) could be activated. Although this must be further explored in terms of intuitiveness and usability in future user-in-the-loop interaction studies, moving away from recognizing a large subset of gestures may enable the robust use of EMG inputs in such consumer use cases. Additionally, it is possible that the input space could be augmented through compound gestures (e.g., two quick swipes), elicitation speed (e.g., slow vs fast swipe), or contraction intensity (i.e., soft and hard double taps). Future work should, therefore, explore how interactive systems can better use smaller sets of EMG-enabled gestures.

### 3.8 Effect of Transfer Learning

#### 3.8.1 Methods

This section evaluates the effect of transfer learning from large cross-user models (i.e., tuning the cross-user LSTM model) with samples from the end user. Two different approaches were explored: a *traditional approach*, and a *contrastive approach*. For both approaches, the three LSTM layers (see Figure 2) were frozen during transfer learning, meaning that only the final linear layers of the network were updated. The two methods, however, differ in their fundamental objective. The traditional approach aimed to tune the network to accommodate the new user’s data, meaning the embedding space could change significantly to accomplish this. The contrastive approach, instead, aimed to project the new user’s data into the original embedding space wherein the original centroids (i.e., anchors) did not change. Both networks were tuned for 400 epochs with an initial learning rate of 1e-2 and a scheduler (*step size=25* and *gamma=0.9*), leading to quick transfer times (*<*1 minute with an NVIDIA RTX 3080). As a baseline for comparison, user-dependent models were also trained as described in Section 2.2. Following the previous exploration in Section 3.6, the effect of transfer learning on model bias was also explored. Finally, the dataset was split into ten repetitions for transfer (or training), 14 repetitions for testing, and a single repetition was held out for validation. The two transfer learning approaches are further described below:

##### Traditional Transfer Learning

This approach updated the final linear layers of the network using cross-entropy loss with the new gesture templates.

##### Contrastive Transfer Learning

This approach aimed to minimize the distance between new samples and the centroids established in the cross-user embedding space by using a loss function loosely inspired from Triplet loss (Hermans et al., 2017), as shown in equation 1. Correspondingly, the final linear layers were tuned to project gestures for new users close to the cross-user centroids in the original embedding space. A k-nearest-neighbour classifier was then trained (where k = the number of repetitions per gesture) to make predictions in the new embedding space (with the new gesture templates).

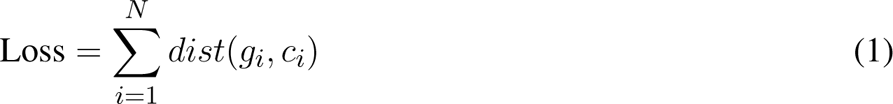

where *i* is the index of the new gesture template, *N* is the number of new gesture templates, *g_i_* is the embedding of the new gesture template, *c_i_* is one of the six centroids from the cross-user embedding space that corresponds to the label of *g_i_*, and *dist* is the euclidean distance.

#### 3.8.2 Results

Figure 11 shows the performance of the two transfer learning approaches. For the user-dependent case (Figure 11A), performance did not reach the zero-shot baseline until seven repetitions were used to train the model. This meant that 3.5 minutes (7 reps x 6 gestures x 5 seconds) of active data collection were required from users before they had performance similar to that of the cross-user model (for which they did not provide any data). While both transfer learning approaches improved the average accuracy (Figure 11B), the contrastive approach generally outperformed the standard approach across all numbers of repetitions. Additionally, both approaches were approaching the 97.9% leave-one-repetition-out user-dependent baseline established in Section 3.1 with half of the needed repetitions. It is also interesting to note that the contrastive transfer approach brings up the absolute bottom end from 43.9% with no transfer to 59.5% with one rep, 70.2% with two reps, and 78.6% with ten reps. This means that the transfer learning approaches were able to help outlier users who had physiological or behavioural differences when eliciting the gestures. Similarly, when applying the contrastive transfer approach, the accuracy of both genders converged at around eight repetitions (Figure 11C). While the accuracies for left and right-handed users did not fully converge (Figure 11D), they became closer. These results indicate that transfer learning approaches can (1) improve the overall accuracy and (2) alleviate some of the effects of biases in the training set.

**Figure 11.**
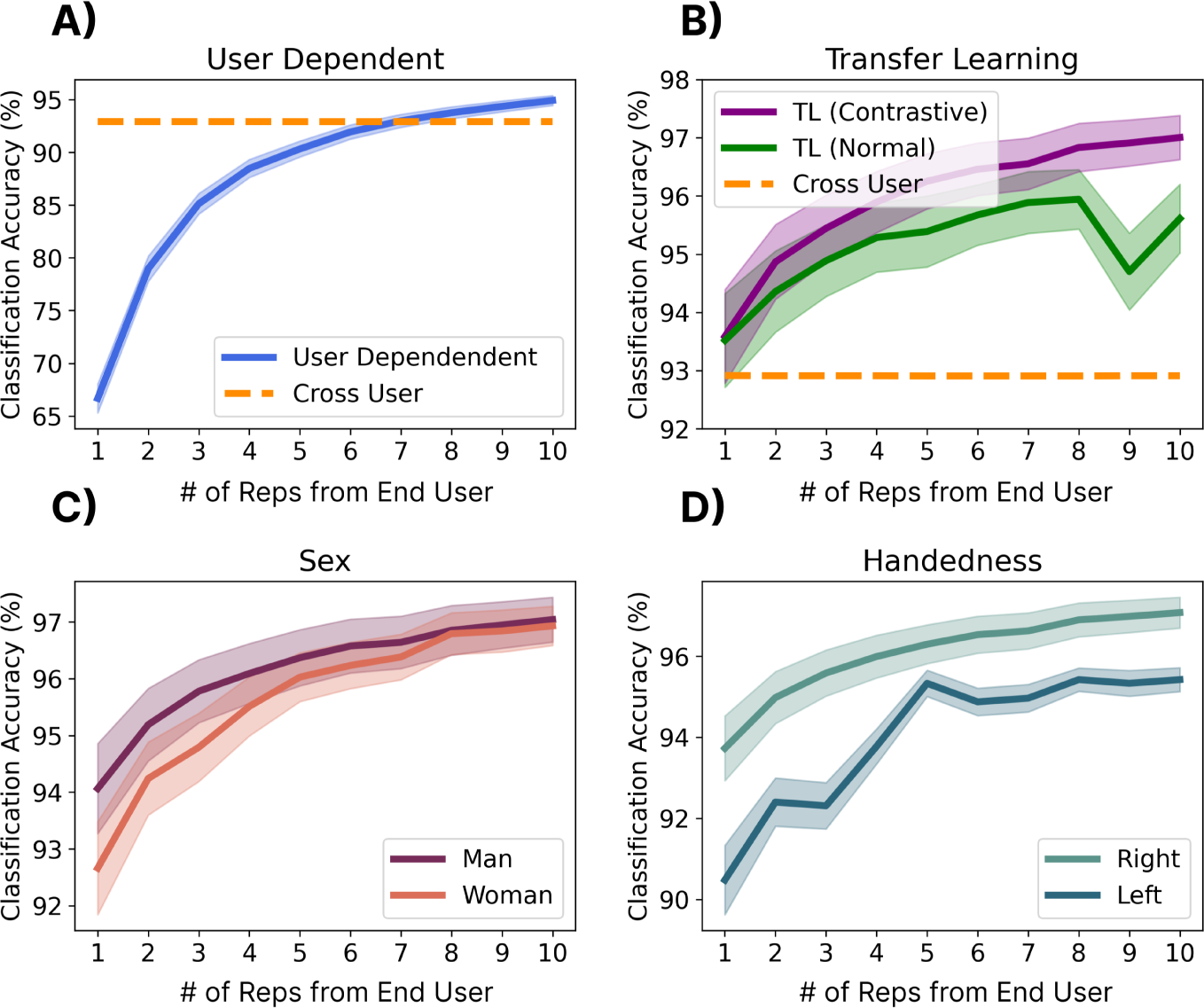
The effect of end-user calibration (i.e., transfer learning) on performance. Shadows indicate standard error. **(A)** Shows the user-dependent performance when training a model from scratch using 1-10 repetitions of each gesture. **(B)** Shows a traditional transfer learning approach where the final linear layers of the cross-user model were tuned to the gestures of the new gesture using Cross Entropy loss and a contrastive transfer learning approach where the loss function minimizes the distance between new gestures and the centroids of the cross-user model. **(C and D)** Show the performance of contrastive transfer learning when splitting results by gender **(C)** and handedness **(D)**.

#### 3.8.3 Discussion

This analysis shows that transfer learning is promising for fine-tuning cross-user models, achieving similar performance to that achieved by the cross-user model in Section 3 with half of the repetitions. It is also possible that, unlike the user-dependent case, the models that have been transferred will be more resilient to confounding factors (see Section 3.9) as these data may have been included in the larger model. The resilience of these transferred models, however, remains to be evaluated in online user-in-the-loop evaluations. One possibility for these approaches would be to leverage them for an initial ‘out-of-the-box’ calibration (similar to the Face ID and fingerprint setup leveraged on mobile devices) or to give users the ability to fine-tune their model (such as with eye calibration for heads-up displays). Alternatively, future work could explore unsupervised adaptive approaches that continuously fine-tune model performance over time without the need for direct calibration (Campbell et al., 2024).

One major benefit of transfer learning was that it improved the bottom end (i.e., outlier participants) by nearly 16%. This could make a substantial difference in equity for individuals who may not be able to use the cross-user model due to factors like limited mobility, thus making these models more accessible. Additionally, these approaches reduced the disparity between underrepresented demographics, including women and left-handed users. These results highlight that even in the case of big-data-enabled zero-shot models, transfer learning may still be beneficial, particularly for improving the model’s accessibility.

### 3.9 Generalization to a New Dataset With Confounding Factors

#### 3.9.1 Methods

This section used the separately recorded dataset described in Section 2.1.2 to evaluate the generalizability of the cross-user model trained on the EMG-EPN612 dataset to other datasets recorded with the same hardware. While this dataset was also recorded with the Myo Armband, the data collection protocols and segmentation strategies employed were different (Eddy et al., 2024b). In particular, this dataset was recorded as part of another study without knowledge of the EMG-EPN612 protocol. This dataset also includes a set of confounding factors that were not intentionally introduced in the EMG-EPN612 dataset to evaluate the robustness of the cross-user model to more representative real-world inputs, including (1) gestures elicited across two different days between one and seven days apart (which necessarily includes removing and re-donning the device), (2) gestures elicited at different speeds (slow, moderate, and fast), and (3) gestures elicited in different limb positions. The limb positions evaluated were: (default) upper arm at side, elbow bent at 90*^◦^* pointed forward along the sagittal axis, (2) upper arm at side, elbow bent at 90*^◦^* pointed away from the body along the frontal axis, (3) upper arm at side, elbow bent at 90*^◦^* in front of the body along the frontal axis, (4) arm fully extended toward the ground, and (5) upper arm relaxed, with elbow full flexed at *∼*180*^◦^*. The Friedman aligned-ranks statistical test with Holm correction was used to check for statistical differences (*p <* 0.05) between the baseline conditions (day 1 and day 2) and the confounds.

#### 3.9.2 Results

The results of the analysis on the separate dataset can be found in Figure 12. The baseline performance of the cross-user model on day one was in line (*∼*93%) with the accuracy achieved using the same model with the held-out testing set from the EMG-EPN612 dataset (see Section 3.2). This implies that the cross-user model was equally robust to data collected from a separate study, in a different country, with different data collection protocols. Interestingly, performance actually improved by nearly 3% on day two, indicating there may have been some learning effect even though no feedback was provided to users. It is possible that, with feedback, these accuracies may continue to improve as the user adapts to the zero-shot model. The results also show that the speed confounding factor significantly degrades (p*<*0.05) classification accuracy, particularly for the slow-speed case. This further reinforces the results from Section 3.6, where outliers on the slower end of the normal distribution curve had the worst performance. Compared to the default speeds elicited on days one and two, the accuracy degraded by 8.4% (*d* = 0.84) and 10.8% (*d* = 1.12), indicating a large effect (d*>*0.8) as measured by Cohen’s d. Finally, the limb position confound was not found to significantly affect classification accuracy except in position 3 (elbow at 90*^◦^* with the hand along the frontal axis) when compared to the day two accuracy. Note that the accuracy in position 3 is still above the 93% cross-user accuracy achieved when using the EMG-EPN612 testing set. However, although no significant difference was found for positions 4 and 5, Figure 12C suggests that some participants may have been affected, as denoted by the wider range of in their lower quartiles.

**Figure 12.**
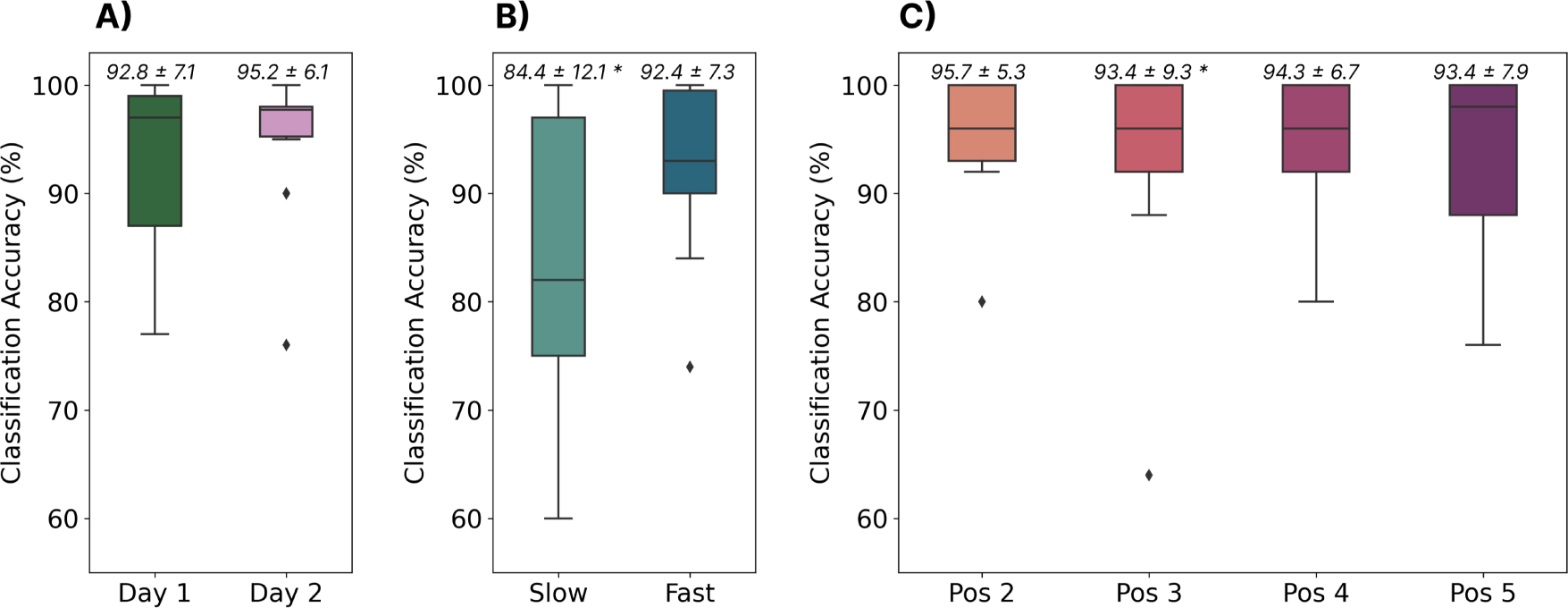
The performance of the cross-user LSTM model when tested using data from a completely different dataset. **(A)** Shows the model’s performance when tested with gestures elicited across two days (the same subjects, between one and seven days apart). **(B)** Shows the performance of the model when tested on gestures elicited at two different speeds (i.e., slow and fast). **(C)** Shows the model’s performance when tested on gestures elicited in four different limb positions. Above each boxplot are the mean and standard deviation where * indicates significant differences (*p <* 0.05) from the day 2 baseline.

#### 3.9.3 Discussion

The results from this analysis highlight two significant findings that may have major implications for the future of myoelectric control. First, large cross-user models can generalize to completely different datasets with different collection protocols and segmentation strategies (recorded nearly five years apart), when the same hardware is used. Future work should explore the ability to transfer pre-trained cross-user models to new hardware devices, as the Myo Armband has since been discontinued. Secondly, the results show that cross-user models can alleviate many of the confounding factors that have historically limited myoelectric control. For example, while previous work has shown that user-dependent models are susceptible to confounds like the limb position effect (Campbell et al., 2020; Fougner et al., 2011; Eddy et al., 2024b), and cross-day use (Jiang et al., 2022; Eddy et al., 2024b), the cross-user model was resilient to both of these factors without them even being intentionally introduced into the dataset. This implies that there was enough variation across the 306 training participants that the model could generalize to these unseen conditions. This further emphasizes that many of the challenges associated with myoelectric control (at least discrete myoelectric control), including usability issues caused by confounding factors, can be alleviated when there is enough data. However, this model was still susceptible to gestures elicited at slower speeds. Correspondingly, future big-data EMG research should consider intentionally introducing some confounds (like speed) into their data collection protocols to further improve the generalizability of these cross-user models. Moreover, future work should evaluate the impact of other confounding factors in the context of cross-user models, such as electrode shift and contraction intensity (Campbell et al., 2020).

### 3.10 Confidence-Based Rejection

#### 3.10.1 Methods

Despite the promising zero-shot cross-user performance seen so far, all tested gestures have been part of a closed-set (consistent with those used to train the model). In many real-world applications, the set of possible gestures will be larger than the set used to train the model, leading to unknown behaviours and erroneous (i.e., false) activations. One often used technique to deal with these so-called out-of-set inputs in myoelectric control is rejection (Robertson et al., 2019; Scheme et al., 2013). Generally, this is done by comparing the confidence or probability output associated with the classifier’s decision with a predefined threshold, below which activation decisions are rejected and ignored. The idea of this approach is that it is better to do nothing (and ignore an input) than to elicit an incorrect action. This could be particularly beneficial in the discrete space, whereby an incorrect decision could result in an action that requires increased time for correction and degrades user confidence in the interface (e.g., a user may accidentally end a phone call or make an erroneous menu selection). Therefore, this section evaluates probability-based rejection for both the within-set case (i.e., rejecting known gestures that may be incorrect due to low certainty) and the out-of-set case (i.e., rejecting gestures that are unknown to the classifier). By using the probability of the classifier (i.e., confidence) associated with each prediction (i.e., in this case, the output from the final softmax layer), the impact of rejection for discrete myoelectric control is explored. The cross-user model was trained as described in Section 3.2, using all of the data from the 306 training subjects. The EMG-EPN612 test set was used to evaluate the closed-set rejection performance. For a given set of rejection thresholds, the percentage of rejected gestures from this closed-set was computed, as well as the new accuracy of the model when omitting these data. The twenty repetitions of each of the gestures that were unique to the Myo DisCo dataset (i.e., index extension, thumbs up, finger snap, and finger gun) were used to evaluate the out-of-set performance of the model.

#### 3.10.2 Results

As shown in the log-scaled Figures 13A and 14A, the confidence profile of the cross-user model was highly skewed towards extremely high confidence for both the within-set and out-of-set gestures. In particular, for the within-set case, 97.3% of gestures were predicted with *>*99.9% confidence. Similarly, for the out-of-set case, 85% of the erroneous gestures were predicted with *>*99.9% confidence. Nevertheless, rejection improved classification accuracy for the within-set case while keeping rejection rates low. For example, using a rejection threshold of 99.99%, the performance of the remaining decisions improved from 92.6% to 94.5% while introducing a rejection rate of only 4.4% (see Figure 13). However, for the out-of-set case, the classifier could only reject 23.0% of the unknown gestures when using the same rejection rate, indicating that dealing with out-of-set inputs still poses a significant challenge for these models.

**Figure 13.**
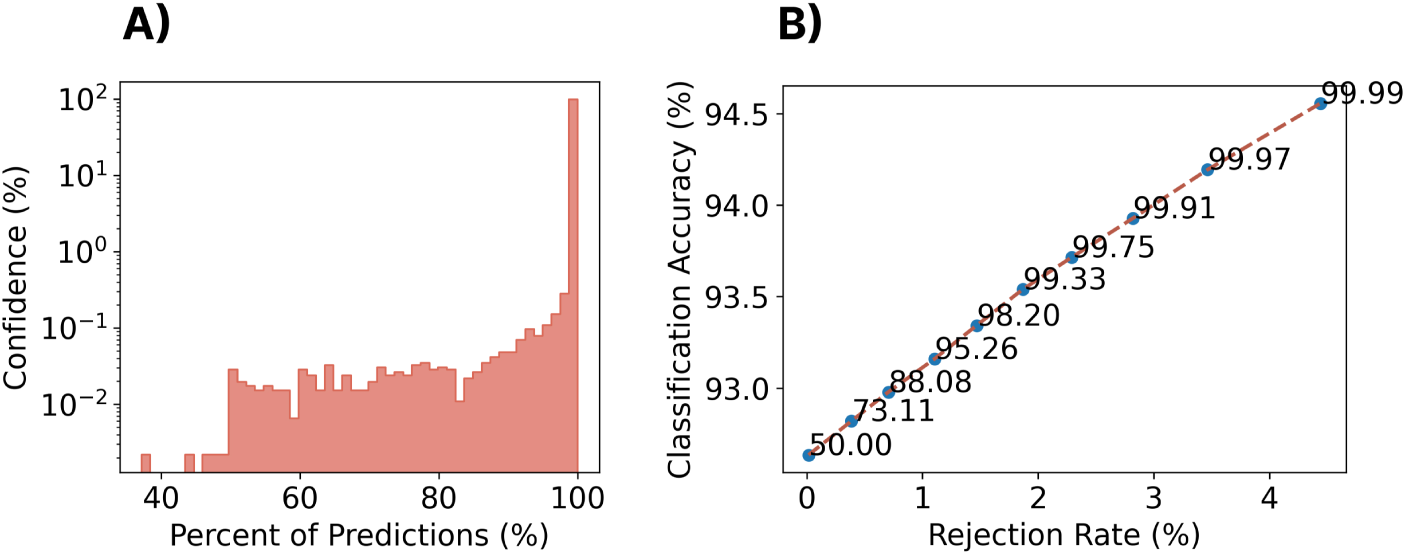
Impact of rejection on the closed-set performance of the zero-shot cross-user LSTM model. **(A)** A histogram of the classifier confidence outputs associated with all testing decisions. *Note that the y-axis is log-scaled.* **(B)** The rejection rate versus classification accuracy for a set of rejection thresholds (each point/label) for the cross-user LSTM model.

**Figure 14.**
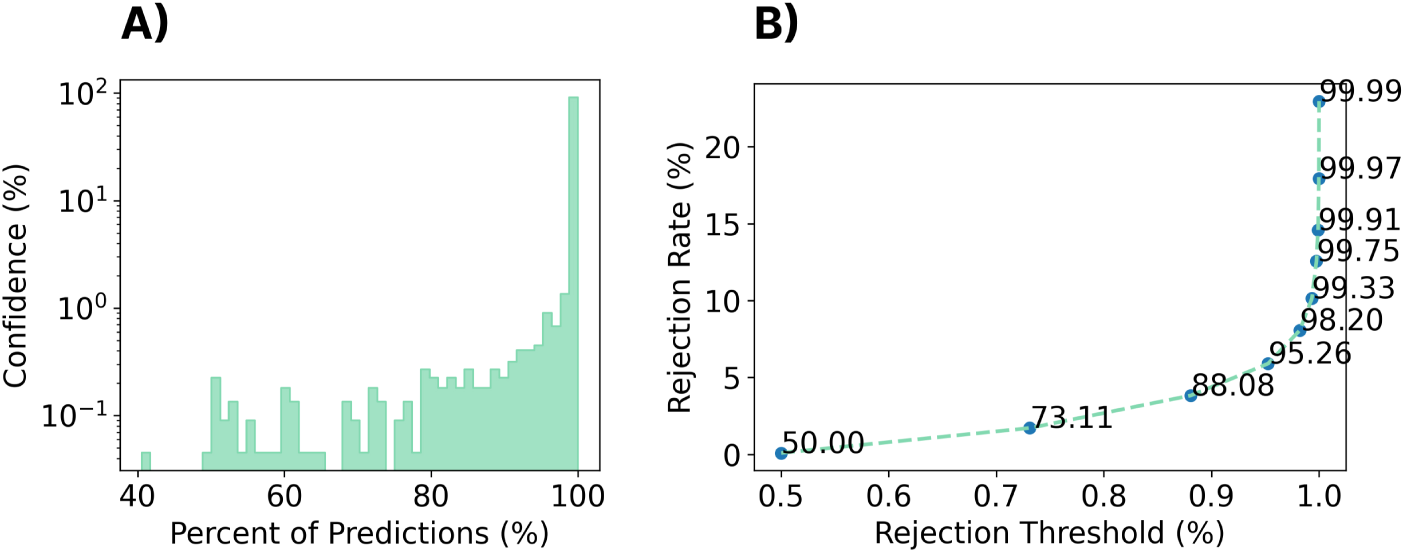
Impact of rejection on unseen, out-of-set gestures when using the zero-shot cross-user LSTM model. The unused gestures from the Myo DisCo dataset (finger gun, finger snap, thumbs up, and point) were used to evaluate the out-of-set performance of the model. **(A)** A histogram of the classifier confidence outputs associated with all out-of-set testing decisions. *Note that the y-axis is log-scaled.* **(C)** The rejection rate of the out-of-set gestures for different rejection thresholds.

#### 3.10.3 Discussion

During the real-world use of a discrete myoelectric control system, users will inevitably make mistakes or elicit inadvertent gestures that could lead to false activations. The results of this preliminary investigation of rejection suggest that even in the case of cross-user models, the confidence distribution of the classifier remains extremely high. Nevertheless, adding a rejection threshold can improve classification performance for the within-set gestures at the expense of a relatively small rejection rate (i.e., rejecting in-set gestures). This tradeoff between true positives and false negatives in online control tasks is an interesting area for future research (Lafreniere et al., 2021).

More importantly, however, the results from this analysis introduce out-of-set rejection as a significant research challenge that still needs to be addressed for discrete myoelectric control. These gestures, along with other patterns of EMG inadvertently generated during activities of daily living (ADLs), will inevitably lead to false activations of system commands. With the current levels of false activation due to out-of-set inputs, an individual out for a run may inadvertently answer an incoming phone call or a teleoperation operator could send an erroneous command to a robot, both of which may have adverse implications. It is worth noting, however, that the out-of-set gestures evaluated in this work were intentionally prompted gestures that were simply not included during training. In real-world applications, patterns of unintended contractions may be more subtle, and by consequence, possibly more rejectable. Furthermore, a user may learn to avoid intentional out-of-set gestures, so further online interaction research is required to fully understand the impact of these false activations. The adoption of highly separable wake gestures (akin to common voice prompts, such as *“Hey Google”*) may also alleviate some of these issues (Kumar et al., 2021; Eddy et al., 2024a), although this may represent a missed opportunity for subtle, always-available ubiquitous input. Even in the recent white paper by Ctrl Labs (Labs et al., 2024), this issue of out-of-set input was not addressed, as all of the control tasks occurred for dedicated tasks. Correspondingly, exploring techniques to improve confidence distributions, such as open set (Bendale and Boult, 2016; Geng et al., 2020) and outlier detection techniques (Zhao et al., 2019), to enable the rejection of incorrect gestures, unknown gestures, and ADLs, is a crucial direction for future research.

## 4 DISCUSSION

### 4.1 Outlook on Zero-Shot Approaches

Despite the physiological and behavioural differences that have historically led to myoelectric model deterioration, this exploration demonstrates that myoelectric gesture recognition can achieve *>*93% performance on unseen users with a large enough dataset. Similar outcomes have recently emerged in other works (Labs et al., 2024), yet this is the first work to comprehensively break down the performance of zero-shot myoelectric gesture recognition models by pattern recognition architecture, training cohort size by both reps and subjects, temporal resolution, feature extraction, and dataset demographics. These outcomes generally support the claim that large datasets are a realistic path to population-level models for myoelectric control of unimpaired subjects.

Moreover, the results show that the zero-shot discrete cross-user model could be generalized to a completely different dataset with a variety of known confounding factors. Not only did the model perform well on this dataset (which was recorded nearly five years later), but the cross-user model effectively reduced and even eliminated the impact of two common confounding factors (limb position variability and cross-day use, respectively). The only caveat was that the dataset was collected with the same type of hardware (*the Myo Armband*), and the armband was oriented in approximately the same configuration for all participants. These results prove that such models can generalize to new use cases and more challenging datasets (Myo DisCo), further highlighting their promise for real-world control. In the future, these pre-trained models could be made publicly available, meaning that anyone with the specified hardware will be able to use them “out-of-the-box”. This could greatly expedite the progression of EMG-based HCI research as authors could easily test against and build off previous research. It is our intention to make these models publicly available in the next major version release of our open-source EMG processing and interaction library, LibEMG (Eddy et al., 2023b).

Despite strong overall performance when averaged across the cohort, imbalances in the EMG-EPN612 dataset led to training biases, resulting in left-handed users, women, and those performing slower gestures performing worse than more represented groups. The exact cause of this performance difference is currently unclear. Is it because insufficient representative information was available from the underrepresented groups? In this case, strategically recruiting these groups to reach satisfactory levels could be a valid solution. Alternatively, is it possible that the abundance of patterns from the normative group overpowered the information from the underrepresented groups? In this case, regularization approaches to retain the information learned for underrepresented groups could be a valid solution. Further, it could be valuable to pinpoint the sources of variability of underrepresented groups through exploratory approaches, like past myoelectric studies using Guided Gradient-weighted Class Activation Mapping (Côté-Allard et al., 2020) or Mapper (Campbell et al., 2019), to avoid relying on information that highlights group differences. Unbalanced datasets are a general issue for deep learning that have led to weaknesses or biases being propagated unknowingly and, as such, should be a subject of future cross-user myoelectric studies.

### 4.2 Outlook on Transfer Learning Approaches

This work has demonstrated zero-shot “plug-and-play” capabilities, where the end-user could hypothetically immediately perform control actions without any additional training protocol; however, algorithmic solutions to personalize to the end-user have historically benefited model performance. A comparison between the user-dependent and user-independent LSTM models still shows a margin of error between these two cases (*∼*98% versus *∼*93%), where a user-dependent model may yield better control in highly constrained use cases. It is conceivable, however, that if such models were used in the real world, the performance would degrade significantly due to confounds such as electrode shift, limb position variability, and gesture elicitation speed (Eddy et al., 2024b), that appear to be alleviated by the variability in the training data of the cross-user models. Furthermore, transfer learning consistently outperformed user-dependent models when the end-user training burden was kept constant (*∼*97% vs.*∼*95% for ten end-user reps). This establishes that transfer learning may be the gold-standard approach for constructing end-user models, especially given that most protocols contain only a few repetitions, aligning with when its benefits are most profound (Phinyomark and Scheme, 2018). Future work should continue to explore the differences between cross-user discrete models with and without transfer in online control tasks.

Although the transfer learning applied in this study was relatively simple, to the author’s knowledge, this was the first study exploring transfer learning, particularly for discrete gesture recognition. Continuous gesture recognition research, however, has explored various algorithmic approaches for transfer learning, namely student-teacher distillation, adaptive normalization approaches, and model search approaches. For instance, a two-stage student-teacher distillation, Dual-Step Domain Adaptation Network (DSDAN), operates by alternating between training a teacher that embeds gestures of many subjects into the same region and distilling teacher-supplied end-user pseudo-labels into a student network, which ultimately adapts to the end-user via an online unsupervised process (Lin et al., 2023). Another approach, adaptive instance normalization, has been used as the fine-tuning approach of the adaptive domain adversarial neural network, whereby the weights of the neural network are shared by all subjects, but the parameters associated with normalization in the network are user-specific. This substitution of normalization parameters allows for the model weights to become user-agnostic during the zero-shot training, and transfer to a new end-user only requires learning their set of normalization parameters using small amounts of supervised data (Campbell et al., 2021). In contrast, supportive convolutional neural networks (Kim et al., 2019) assume a large number of pre-trained classifiers are available of which a subset should provide reasonable zero-shot performance on a trial of end-user data. From this subset, independent fine-tuning of weights is performed with the same trial of end-user data, after which the mode of these fine-tuned classifiers are used for subsequent predictions. Further work is warranted, looking into the efficacy of such transfer learning approaches on discrete gesture recognition with large datasets.

### 4.3 Outlook on Online Discrete Gesture Recognition

Historically, EMG researchers have acknowledged that there is only a moderate correlation between offline metrics (like classification accuracy) and user-in-the-loop continuous myoelectric control (Lock et al., 2005; Campbell et al., 2024; Nawfel et al., 2021). Ultimately, many varying user behaviours (e.g., modulation of contractions to get the desired output) result in vastly different patterns than the relatively static and consistent gestures collected during screen-guided training. Additionally, these systems are quite susceptible to confounding factors as they are often highly tuned to recognize specific characteristics from individual windows of data (Scheme and Englehart, 2011). When the signal characteristics change even slightly (e.g., the armband is slightly shifted or a source of noise is introduced), large errors begin to arise. As a result, continuous gesture recognition benefits greatly from obtaining patterns produced from user-in-the-loop settings for adaptation (Woodward and Hargrove, 2019), which could be seen as another fine-tuning procedure beyond collecting end-user data through screen-guided training. This means that, at the moment, continuous myoelectric control has an extensive training burden associated with achieving robust online performance which has hindered the perception of EMG-based inputs (Campbell et al., 2024; Szymaniak et al., 2022; Huang et al., 2024).

Compared to continuous control, however, it is likely that offline metrics like classification accuracy will be more tied to online performance for event-driven discrete myoelectric control systems such as those explored in this work. This is partly because discrete inputs do not allow users to modulate their contractions until after the entirety of a gesture has been elicited. This means that users can only moderate their gestures after feedback at the end of the gesture (i.e., the event was or was not triggered), leading to potentially less varied behaviours. Moreover, when recognizing an entire gesture template consisting of multiple windows, there becomes more data to make informed decisions than in the continuous control case. These additional data, combined with an imposed temporal structure, may be responsible for the high recognition accuracies achieved in this work. Nevertheless, it is crucial that the cross-user models developed here (and any in the future) be tested in online control tasks. This becomes particularly interesting for cross-user models as users will tend to modulate their behaviours to accommodate the model rather than the model accommodating the user’s behaviours (as is common for user-dependent myoelectric control). Additionally, future work should directly explore the correlation between offline metrics and user-in-the-loop control for discrete myoelectric control systems. For example, users may rush gestures in contexts that promote urgency (e.g. cancelling an alarm in public), leading to pattern changes. Finally, a factor that may hinder online control for discrete systems that has not been an issue for continuous control of prostheses is the presence of activities of daily living (patterns of contractions elicited by users that unintentionally resemble target gestures while doing other activities) (Chang et al., 2020). As highlighted in this work, the cross-user models are still quite susceptible to false activations when presented with out-of-set gestures, which would inhibit a user’s ability to engage in other tasks. To improve the usability of discrete systems, future work should thus explore techniques to enable discrete myoelectric control in everyday settings where inadvertent (non-intentional) system inputs will be inevitable.

### 4.4 A Coming Together for Recording Large Datasets

Although much progress has been made for EMG-based control over the previous decades, there remains an ongoing challenge limiting the true progression of the technology: a lack of large multi-user datasets. Due to non-standardized hardware and data collection protocols, many researchers endeavour to collect study-specific datasets that either never get publicly released or cannot be pooled with other pre-recorded data. In particular, other than the 612 user dataset used in this study (Benalcazar et al., 2021), and the private collection done by Ctrl Labs (Labs et al., 2024), very few large (*>*100 user) EMG datasets have been recorded, and almost none are publicly available. This is contrary to other machine learning fields, whereby large datasets have become commonplace. For example, ImageNet contains 3.2 million images (Deng et al., 2009), People’s Speech includes 30,000 hours of annotated speech data (Galvez et al., 2021), and the MNIST dataset contains 70,000 annotated images of handwritten digits (Deng, 2012). These large open-source datasets can be attributed to the rapid advancement of computer vision and speech recognition, which are both now commercialized in many devices we use daily.

Moving forward, the EMG community must begin finding ways to curate large datasets, as has been done in other machine learning communities. One technique to enable this is publishing data collection protocols whereby authors can contribute to larger data repositories. Open-source tools like LibEMG or new platforms to enable EMG data collection with standardized protocols could facilitate this process (Eddy et al., 2023b). Alternatively, a potential research direction could be to find ways to allow the pooling of data from different collection protocols and hardware. Regardless, as a community, we must come together to create these large datasets as they could have significant benefits beyond general-purpose EMG, potentially also improving the robustness of myoelectric control for powered prosthetics, and more.

## 5 CONCLUSION

For a long time, myoelectric control has been fraught with challenges that have limited its use outside of powered prostheses. This work shows that many of these issues, including the inter-subject differences and susceptibility to confounding factors like limb position variability and cross-day use, can largely be eliminated when training large multi-user models for discrete control (*>*300 users). In particular, the results suggest that zero-shot discrete cross-user models can achieve classification accuracies of up to *∼*93% for recognizing six discrete gestures from a set of 306 unseen testing users. While these results highlight the potential promise of myoelectric control as a general-purpose ubiquitous input, this work highlights many future challenges and research directions that must be explored before its eventual success and widespread adoption.

## AUTHOR CONTRIBUTIONS

**Ethan Eddy** Conceptualization, Formal Analysis, Investigation, Methodology, Software, Writing - Original Draft. **Evan Campbell** Conceptualization, Writing - Review and Editing. **Scott Bateman** Conceptualization, Supervision, Writing - Review and Editing. **Erik Scheme** Conceptualization, Methodology, Supervision, Writing - Review and Editing.

## ACKNOWLEDGMENTS

The authors would like to acknowledge the important contributions of the authors of the EMG-EPN612 dataset (Benalcazar et al., 2021); Marco E. Benalcazar, Lorena Barona, Leonardo Valdivieso, Xavier Aguas, and Jonathan Zea, without which this work would not have been possible.

## DATA AVAILABILITY STATEMENT

The primary dataset used for this study can be found in the EMG-EPN612 repository: https://laboratorio-ia.epn.edu.ec/es/recursos/dataset/2020_emg_dataset_612.

## Footnotes

1 https://www.apple.com/watch/

